# Photonic chip-based multimodal super-resolution microscopy for histopathological assessment of cryopreserved tissue sections

**DOI:** 10.1101/2021.05.06.442952

**Authors:** Luis E. Villegas-Hernández, Vishesh Dubey, Mona Nystad, Jean-Claude Tinguely, David A. Coucheron, Firehun T. Dullo, Anish Priyadarshi, Sebastian Acuña, Jose M. Mateos, Gery Barmettler, Urs Ziegler, Aud-Malin Karlsson Hovd, Kristin Andreassen Fenton, Ganesh Acharya, Krishna Agarwal, Balpreet Singh Ahluwalia

## Abstract

Histopathological assessment involves the identification of anatomical variations in tissues that are associated with diseases. While diffraction-limited optical microscopes assist in the diagnosis of a wide variety of pathologies, their resolving capabilities are insufficient to visualize some anomalies at subcellular level. Although a novel set of super-resolution optical microscopy techniques can fulfill the resolution demands in such cases, the system complexity, high operating cost, lack of multimodality, and low-throughput imaging of these methods limit their wide adoption in clinical settings. In this study, we interrogate the photonic chip as an attractive high-throughput super-resolution microscopy platform for histopathology. Using cryopreserved ultrathin tissue sections of human placenta, mouse kidney, and zebrafish eye retina prepared by the Tokuyasu method, we validate the photonic chip as a multi-modal imaging tool for histo-anatomical analysis. We demonstrate that photonic-chip platform can deliver multi-modal imaging capabilities such as total internal reflection fluorescence microscopy, intensity fluctuation-based optical nanoscopy, single-molecule localization microscopy, and correlative light-electron microscopy. Our results demonstrate that the photonic chip-based super-resolution microscopy platform has the potential to deliver high-throughput multimodal histopathological analysis of cryopreserved tissue samples.

## 2. Introduction

Histopathology refers to the study of tissue sections under a microscope to diagnose diseases, guide medical treatment, and prognose clinical outcomes. To date, this well-established discipline is one of the key decision-support tools available for clinicians across the world. A typical histological analysis involves the extraction of a tissue sample from the body, fixation, and preservation followed by sectioning, labeling, and microscopy. By performing a morphological assessment of the tissue under the microscope, histopathologists can identify various diseases and render a clinical diagnosis.

Imaging throughput, contrast, and resolution are critical parameters in the histopathological assessment. Given the morphological heterogeneity of the samples, pathologists often need to assess tens to hundreds of cm^2^ section areas to locate and analyze the lesions [1]. Thus, high-throughput imaging platforms are desirable for routine histopathological analysis. The whole slide imaging scanners fulfill this requirement by allowing fast imaging of several histological slides in a day. However, these automated optical microscopes are limited to a resolution power of ~250 - 500 nm [2], which is insufficient for the identification of some pathologies, for example, nephrotic syndrome and amyloidosis. For decades, the visualization of such pathologies was only possible through other imaging techniques such as electron microscopy [3, 4], which supports a resolving power down to ~10 nm for fixed and embedded histological samples. However, the combination of a lengthy sample preparation process, a low imaging throughput, and the lack of specificity makes electron microscopy an inconvenient and costly technique for clinical use, hindering its broad adoption for routine histopathological examination of tissue samples and restricting its implementation to basic-biology research.

Recently, the advent of super-resolution fluorescence optical microscopy techniques, also referred to as optical nanoscopy, bridged the resolution gap between the diffraction-limited optical microscopy and the electron microscopy methods, allowing for high-specificity imaging of biological specimens at high-resolution [5–7]. Present-day fluorescence-based super-resolution optical microscopy comprises a panel of methods that exploit engineered illumination and/or the photochemical and photokinetic properties of fluorescent markers to achieve high spatio-temporal resolution. These include structured illumination microscopy (SIM), stimulated emission depletion microscopy (STED), singl-molecule localization microscopy (SMLM), and intensity fluctuation-based optical nanoscopy techniques (IFON).

While super-resolution fluorescence optical microscopy methods are commonly used in cell biology, their adoption in histopathological settings remains deferred due to multiple reasons: a) the high labelling density of tissues poses challenges on super-resolution methods, especially for SMLM and IFON, where a high spatio-temporal sparsity is necessary for the reconstruction of structures beyond the diffraction limit of the microscope; b) the susceptibility of the super-resolution methods to optical aberrations and light scattering introduced by refractive index variations across the samples [8]; c) the imaging artifacts induced by autofluorescence signal of tissues [9]; and importantly, d) the low throughput, high-cost, lack of multi-modality, system complexity, and bulkiness of existing super-resolution optical microscopy setups.

Although, limited work on using STED, SIM, and SMLM have been explored for super-resolution imaging of tissues [10–12], these methods fail to fulfill the throughput demands for routine histopathological assessment (see *Supplementary Information S1*). For example, STED, albeit delivering a lateral resolution down to 20 nm [13] and being a robust confocal method to scattering challenges posed by tissues, is an inherently low-throughput point-scanning technique. Similarly, SMLM and SIM, despite being wide-field methods supporting sub-50 nm and ~110 - 130 nm lateral resolution respectively, are heavily dependent on the acquisition of multiple frames and subsequent reconstruction via post-processing algorithms. While SIM outperforms SMLM in terms of imaging speed, requiring only 9 or 15 images (2D/3D cases accordingly) as compared to the tens of thousands of images necessary for SMLM, the field of view obtained by commercial SIM systems is typically limited to about 40 × 40 μm^2^. Importantly, among all the super-resolution methods, SIM has been proposed for high throughput imaging in histopathological settings [1, 11]. However, these approaches focused on the acquisition of large field of view images using low magnification and low numerical aperture objective lenses, compromising the lateral resolution to a maximum of 1.3 μm. In terms of system complexity, SMLM is simpler to implement as compared to SIM and STED, which require more sophisticated, bulkier, and costly setups. From an overall perspective, improvements in imaging throughput and reductions in system complexity, footprint, and cost are needed for the adoption of super-resolution fluorescence optical microscopy in histopathology. It is evident from *Supplementary Table S1* that different imaging methods offer different technical capabilities. Thus, to enable widescale penetration in the clinical settings, it is desirable to have an imaging platform that can deliver different super-resolution capabilities using standard optical microscopy setup.

Another important aspect for the adoption of fluorescence-based super-resolution optical microscopy is the availability of a large selection of fluorophores. While SIM works with photo-stable and bright fluorophores, STED and SMLM are more restricted to a special type of fluorescent markers. Interestingly, some of the IFON techniques, such as the multiple signal classification algorithm (MUSICAL) [14], can exploit the pixel intensity variations arising not only from the intrinsic fluctuations of the fluorophores but also from the modulated emissions generated via engineered illumination, enabling a practical implementation with almost all kinds of fluorophores. Despite being an attractive route to follow for clinical applications in histopathology, to the best of our knowledge the engineered illumination approach for IFON has not been explored in tissue imaging.

In recent years, photonic chip-based nanoscopy has emerged as a promising imaging platform for biological applications [15–18], supporting high-resolution, high-throughput, and multi-modal capabilities. To date, photonic chip-based microscopy studies have focused primarily on cellular biology [15–21], leaving on-chip histological imaging relatively unexplored. In this work, we interrogate the photonic chip-based imaging platform to address some of the challenges related to super-resolution imaging of tissue sections. We start by evaluating the viability of the photonic chip for diffraction-limited total internal reflection microscopy (chip-TIRFM). Then, we transition to more advanced chip-TIRFM based imaging methods such as SMLM, IFON, to conclude with a correlative light-electron microscopy (CLEM) analysis. Among the existing histological methods, we chose the *Tokuyasu* protocol [22] for the preparation of the tissue sections. This cryosectioning method provides excellent ultrastructural preservation, high molecular antigenicity, and a thin section thickness (70 nm to 1 μm) that assists both in reducing the light scattering artifacts associated with thicker samples [23] and in making optimal use of the illumination delivered by the photonic chip. We describe the staining protocols and the imaging parameters necessary for photonic chip-based microscopy of tissue samples and discuss the challenges and the advantages offered by this imaging platform for histopathology. By exploiting the engineered illumination delivered by the photonic chip-based microscopy, we further demonstrate the suitability of this novel technique as a compact, high-resolution, high-contrast, high-throughput, and multi-modal imaging platform for histopathology.

## 3. Photonic chip-based microscopy for histopathology

In chip-based microscopy, a photonic chip is used both to hold the sample and to provide the excitation illumination necessary for fluorescent emission (Figure 1), while a standard upright microscope is used to acquire the image (Figure 1c). The photonic chip is composed of two substrate layers of silicon (Si) and silicon dioxide (SiO_2_), respectively, and a biocompatible waveguide core layer that transmits visible light, made of either silicon nitride (Si_3_N_4_) [17] or tantalum pentoxide (Ta_2_O_5_) [24]. Upon coupling, the excitation laser beam is tightly confined inside the optical waveguide layer and propagates through its geometry via total internal reflection (Figure 1a). This generates an evanescent field on the top of the waveguide surface with a penetration depth of up to ~150 - 200 nm that is used to excite the fluorescent markers located in the vicinity of the waveguide surface (see *Supplementary Information S2*). The fluorescent emission is then collected by a standard microscope objective, enabling chip-TIRFM (Figure 1e).

**Figure 1.**
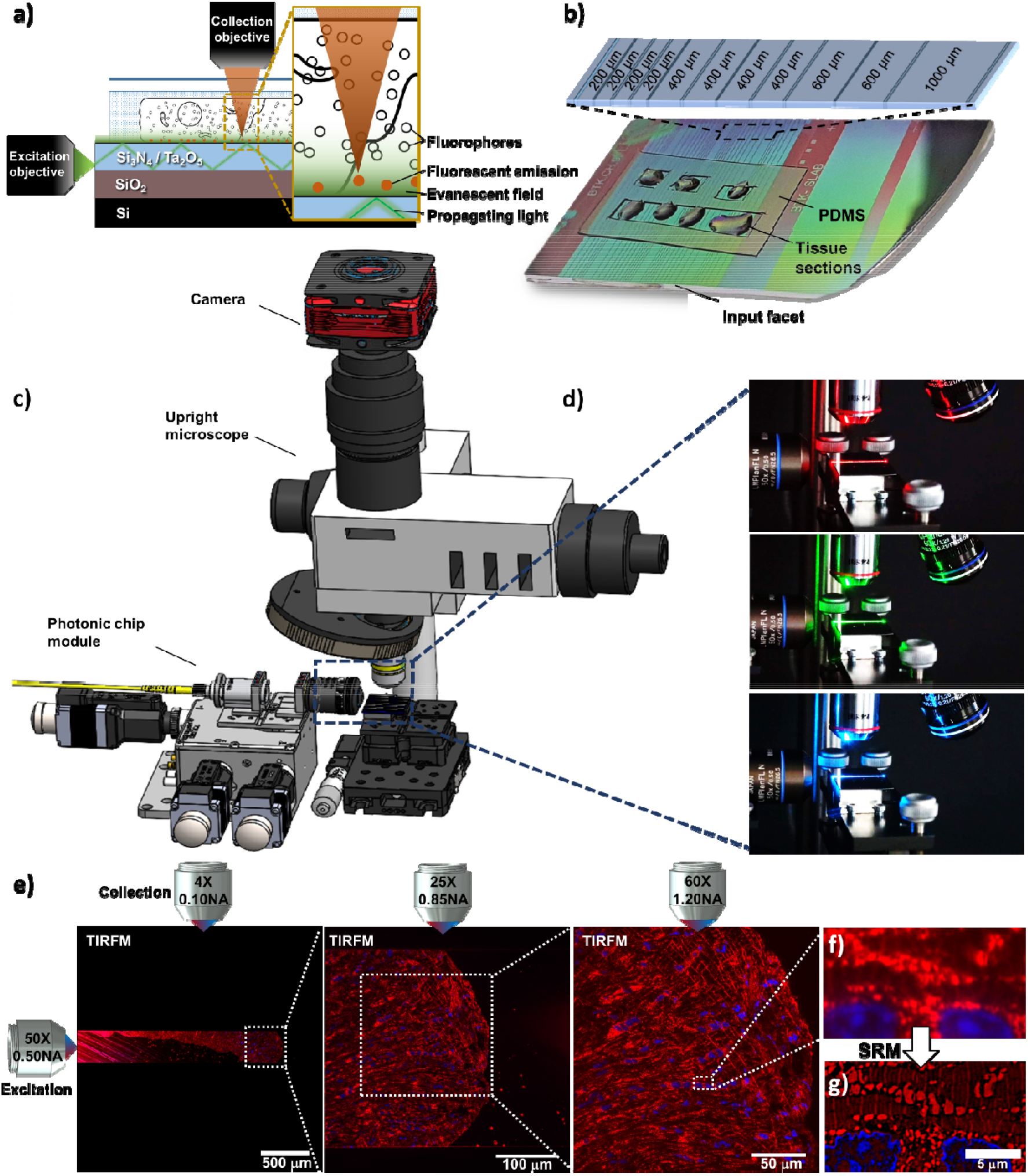
Schematic representation of the chip-based total internal reflection fluorescence microscopy (chip-TIRFM) setup. **(a)** Working principle of chip-TIRFM: upon coupling onto the input facet, the excitation light propagates through the waveguide core material due to total internal reflection. An evanescent field of approx. 150 nm height excites a thin layer of fluorescent dyes in the vicinity of the photonic chip surface, allowing for TIRFM imaging. **(b)** Top view of a photonic chip containing ultrathin Tokuyasu cryosections covered with a 1:1 cryoprotectant mixture of 2.3M sucrose and 2% methylcellulose, and surrounded by a custom-made transparent polydimethylsiloxane (PDMS) frame. The inset illustrates the various strip waveguide widths available on the chip. **(c)** The chip-TIRFM setup is composed of a custom-made photonic chip module and a commercially available upright collection module. Upon coupling the excitation light on the photonic chip, the fluorescent signal is allowed through a filter set and captured with a scientific CMOS camera. **(d)** The photonic chip allows decoupling of the excitation and the collection light paths, enabling TIRFM imaging using conventional microscope objectives. Different wavelengths propagating on the waveguide core allow for multicolor TIRFM imaging. **(e)** TIRFM images of a 100 nm thick pig heart cryosection imaged on a photonic chip through different microscope objectives. Membranes in red and nuclei in blue. **(f)** Magnified view of the diffraction-limited TIRFM image acquired with a 60X/1.20NA water immersion microscope objective. **(g)** Subsequent post-processing of the raw data enables super-resolution microscopy (SRM), allowing the visualization of structures beyond the diffraction limit of conventional optical microscopy.

Photonic chip-based illumination provides several advantages that can be exploited for super-resolution imaging of histopathology samples such as:

a. The photonic chip allows decoupling of the excitation and the emission light paths, which translates into high-contrast images with improved imaging throughput. The propagating light enables a uniform illumination over the entire length of the waveguide while providing optical sectioning of the sample via evanescent field excitation [25]. As the illumination is provided by the photonic chip, the imaging objective lens can be freely changed (Figure 1d), enabling the acquisition of images over large fields of view [16] (Figure 1e), a feature not available in conventional TIRFM setups.
b. The multi-mode interference illumination generated on the photonic chip assists in generating the necessary emission sparsity for diverse super-resolution fluorescent optical microscopy methods, as recently demonstrated via on-chip IFON [15, 26], on-chip SMLM [15, 16, 27], and on-chip SIM [28]. Moreover, by using waveguide materials of high refractive index (for example, *n* = 2), it is possible both to tightly confine the light and to generate higher spatial frequencies as compared to free-space optical components [15, 28], which can be further exploited by IFON techniques such as MUSICAL to super-resolve highly dense and heterogeneous samples such as tissues.
c. Correlative imaging with other established methods including electron microscopy [29] and quantitative phase microscopy [24] can be seamlessly implemented on the photonic chip, expanding the opportunities both for routine analysis and for basic histopathology research.
d. The photonic chip-based microscopy can be implemented on standard optical microscopy platforms upon few adaptations for the integration of a photonic chip module (Figure 1c). The photonic chips can be manufactured in high-volumes following standard complementary metaloxide-semiconductor (CMOS) photolithography processes, allowing for low operating costs in clinical settings.

## 4. Results and discussion

### 4.1 Chip-based multicolor TIRFM imaging

In this part of the study, we used chorionic villi tissue from human placenta to assess the suitability of the photonic chip for histological observations. This tissue, present on the fetal side of the placenta, is responsible for the air, nutrient, and waste exchange between the mother and the fetus during pregnancy [30], and is characterized by villous-like structures, namely villi, that sprout from the chorionic plate of the placenta to maximize the maternofetal transfer processes and communication. When transversally sectioned, the chorionic villi appear in the form of rounded islands distributed across an open space surrounded by maternal blood, called the intervillous space.

Developed by Kiyoteru Tokuyasu in the’70s [31, 32], the so-called “Tokuyasu method” is still a gold standard protocol for ultrastructural analysis of cells and tissues [22]. Primarily established for EM techniques, recent studies have shown its versatility in fluorescence microscopy [33, 34]. For chip-based multicolor TIRFM imaging, 400 nm thick chorionic villi cryosections were prepared following the Tokuyasu method (see detailed preparation protocol in Materials and Methods and *Supplementary Information S3*). After cutting the tissue blocks on a cryo-ultramicrotome, the sections were deposited onto a photonic chip previously coated with poly-L-lysine and equipped with a custom-made transparent polydimethylsiloxane (PDMS) frame (Figure 1b). The membranes, F-actin, and nuclei were fluorescently labeled using CellMask Deep Red, Phalloidin-Atto565, and Sytox Green, respectively.

For the excitation of the respective fluorescent dyes, three independent laser light wavelengths were used, namely 640 nm, 561 nm, and 488 nm (Figure 1d). To obtain TIRF images (see detailed acquisition steps in *Materials and Methods*), the excitation light was coupled onto a single strip waveguide using a 50X/0.5NA microscope objective (Figure 1c). Upon coupling, a multi-mode interference pattern was generated along the waveguide by the propagating light, which could be modulated by changing the position of the coupling objective relative to the chip (see *Supplementary Information S4*). To deliver a uniform illumination onto the sample, the coupling objective was laterally scanned along the input facet of the chip while individual frames were acquired. The fluorescent emission was collected by standard microscope objectives transitioning from lower to higher magnification to achieve different fields of view. Thereafter, the collected signal was averaged, pseudo-colored (membranes in red, F-actin in green, and nuclei in blue), and merged, allowing multicolor visualization of the different tissue components.

The large field of view provided by the 4X/0.1NA objective lens enabled us to locate the sample on the waveguide (Figure 2a), while the 20X/0.45NA assisted for a contextual visualization of the tissue structure, supporting the identification of regions of interest for imaging with further magnification (white box in Figure 2b). Finally, with the aid of a 60X/1.2NA water immersion objective lens (Figure 2c), it was possible to visualize relevant structures of the chorionic villi, such as the apical layer of syncytiotrophoblastic cells, and the abundant fetal capillaries. Arguably, in this study, the absence of maternal red blood cells in the intervillous space can be attributed to the rinsing steps carried out along with the sample collection (see *Materials and Methods*). Figure 2c also allows the visualization of multinucleated cell aggregates that resemble the syncytial knots usually deported onto the maternal blood at different stages of the pregnancy [35]. Notably, the membrane marker not only allowed for an overall view of the tissue (Figure 2b,c) but also enabled the distinction between adjacent cells such as a cytotrophoblast cell and a syncytiotrophoblast cell (white box in Figure 2c and magnified view in Figure 2d). Moreover, the observed F-actin signal (Figure 2e) matched the locations reported in a previous study [36], allowing the identification of the microvilli brush border, the syncytiotrophoblastic’s basal cell surface, and the capillary endothelial cells.

**Figure 2.**
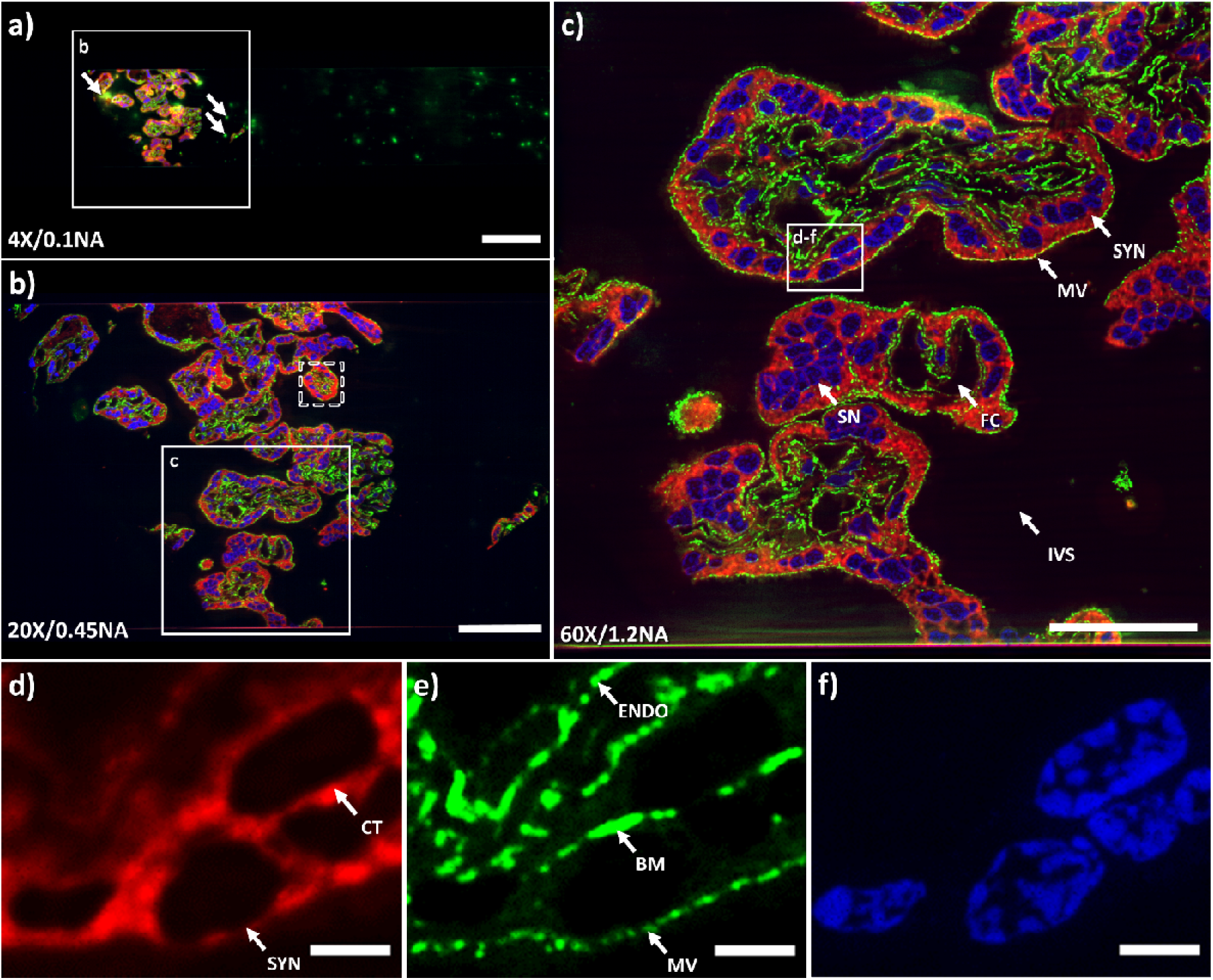
Chip-based multicolor TIRFM imaging of a 400 nm placental tissue section prepared by Tokuyasu method. Membranes labeled with CellMask Deep Red (pseudo-colored in red), F-actin labeled with Phalloidin-Atto565 (pseudo-colored in green), and nuclei labeled with Sytox Green (pseudo-colored in blue). **(a)** Large field of view chip-based multicolor TIRFM image acquired with a 4X/0.1NA microscope objective. The white arrows indicate the locations of unspecific binding of the F-actin marker to the waveguide. The white box represents the area imaged with a higher magnification objective lens in (b). **(b)** Chip-based multicolor TIRFM image acquired with a 20X/0.45NA microscope objective. The white box represents the area subsequently imaged with a higher magnification objective lens in (c). The white-dotted box illustrates the maximum field of view (50 μm × 50 μm) attainable in a conventional TIRFM setup. **(c)** Multicolor chip-TIRFM image acquired with a 60X/1.2NA microscope objective allows the identification of morphologically relevant structures of the chorionic villi such as the syncytiotrophoblastic cells (SYN), fetal capillaries (FC), syncytial knots (SN), and intervillous space (IVS) without maternal red blood cells due to thorough rinsing during sample preparation. The white box corresponds to the individual channels magnified in (d-f). **(d)** A magnified view of the membrane signal allows the distinction between a SYN and a cytotrophoblastic cell (CT). **(e)** A magnified view of the F-actin signal conforms to the expected location for this marker, in places such as the microvilli brush border (MV), the SYN’s basal membrane (BM), and the capillary endothelial cell (ENDO). **(f)** Magnified view of syncytial and cytotrophoblast nuclei. Scale bars (a) 200 μm, (b) 100 μm, (c) 50 μm, (d-e) 5 μm.

The cross-sectional dimensions of the Tokuyasu sections (typically ranging between 300 × 300 μm^2^ and 500 × 500 μm^2^) perfectly suited the waveguide dimensions of the photonic chip used in this work. This configuration allows both complete imaging of the sample through a single optical waveguide and also supports independent illumination of adjacent waveguides on the chip with different tissue sections (Figure 1b). This eliminates undesired excitation light of the samples outside the imaging region of interest, hence minimizing photobleaching. Moreover, the PDMS chambers (Figure 1b) allowed multi-well experiments similarly to traditional microscope chamber slides, with the additional advantage of reducing the incubation volumes to approximately 10 μl to 20 μl per chamber, which translated into a cost-reduction of the fluorescence assays. After optimizing the sample preparation and imaging steps (see *Supplementary Information S5, S6, and S7*), we were able to both fluorescently label and acquire chip-TIRFM images of placental tissue within a timeframe of three hours from cryosectioning to image post-processing.

For diffraction-limited imaging of tissue samples, such as shown in Figure 2, the evanescent field illumination supported by TIRFM is not necessary. However, for super-resolution methods such as SMLM and IFON, the evanescent illumination generated by the photonic chip configuration plays a key role in supporting optical sectioning of the specimen, reducing the out-of-focus light, increasing the signal-to-background ratio, and improving the axial resolution. Conventional TIRFM setups use oil-immersion high numerical aperture (N.A. 1.47-1.50) and high-magnification objective lenses (60X - 100X) that limit their field of view to around 50 × 50 μm^2^ [20] (dotted box in Figure 2b). On contrary, the photonic chip-based TIRFM setup allows the use of essentially any imaging objective lens for the collection of the fluorescent signal, achieving scalable resolution and magnification on demand and opening possibilities for large TIRFM imaging areas up to the mm^2^ scale (Figure 2a, b). To this extend, the photonic chip-based TIRFM technique has the potential to outperform traditional ways of generating an evanescent field, which can be exploited for super-resolution imaging, as detailed in the next sections.

### 4.2 Chip-based SMLM imaging

In the previous section, we demonstrated the suitability of the photonic chip for diffraction-limited TIRFM imaging of tissues. Here, we explored on-chip super-resolution imaging of tissue samples using single-molecule localization microscopy (SMLM) [5]. SMLM comprises a set of methods that exploit the stochastic activation of individual fluorescent molecules to enable their precise localization within a sub-diffraction limited region. To achieve this, the fluorescent molecules are manipulated to obtain sparse blinking events over time. In practice, the majority of the fluorophores are switched off (not emitting light), while only a small segment of them is switched on (emitting fluorescence). This implies the collection of several thousands of frames for the localization of the individual molecules in the sample.

There exist multiple variants of SMLM employing diverse switching mechanisms. Among them, the *direct* stochastic optical reconstruction microscopy (*d*STORM) method supports conventional fluorophores, delivers a high photo-switching rate, and offers low photobleaching [37]. To explore the capabilities of the photonic chip for SMLM on histological samples, we used a 400 nm thick mouse kidney cryosection. We employed a *d*STORM approach to visualize the ultrastructural morphology of the filtration compartments present in the renal tissue, called glomeruli, whose physical dimensions are typically beyond the resolution limit of conventional optical microscopy and, therefore, often studied through electron microscopy.

The membranes and the nuclei were fluorescently stained with CellMask Deep Red and Sytox Green, respectively. All the preparation steps were performed identically to the *chip-based multicolor TIRFM imaging experiments*, except for the mounting medium that consisted of a water-based enzymatic oxygen scavenging system buffer [15, 16] (see details in *Supplementary Information S8*). This oxygen scavenging buffer induces the blinking behavior by enhancing the probability of the fluorescent molecules to transition into the dark state, thereby contributing to the temporal sparsity of emission necessary for SMLM.

To find the features of interest, a TIRFM image of the sample was acquired (Figure 3a) using low laser power to avoid photo-switching and reduce the chances of photo-bleaching. Next, the laser power was increased until sparse blinking was observed. The camera exposure time was set to around 30 ms to capture individual emission events of the membrane dye while the coupling objective was randomly scanned along the input facet of the chip. The collected image stack (> 40,000 frames) was computationally processed to localize the spatial coordinates of the fluorophores, allowing for the reconstruction of a super-resolved image (Figure 3b). A comparative view of both methods (Figure 3c) reveals structural details in *d*STORM that are not discernible in diffraction-limited TIRFM. In particular, *d*STORM allows the visualization of a ~100 nm gap between the podocytes and the endothelial cells (see the empty gap between the white arrows in Figure 3c), which is in agreement with the morphology of the glomerular basal membrane [38]. The identification of this feature, in particular, may be of critical value for a faster diagnosis of nephrotic diseases.

**Figure 3.**
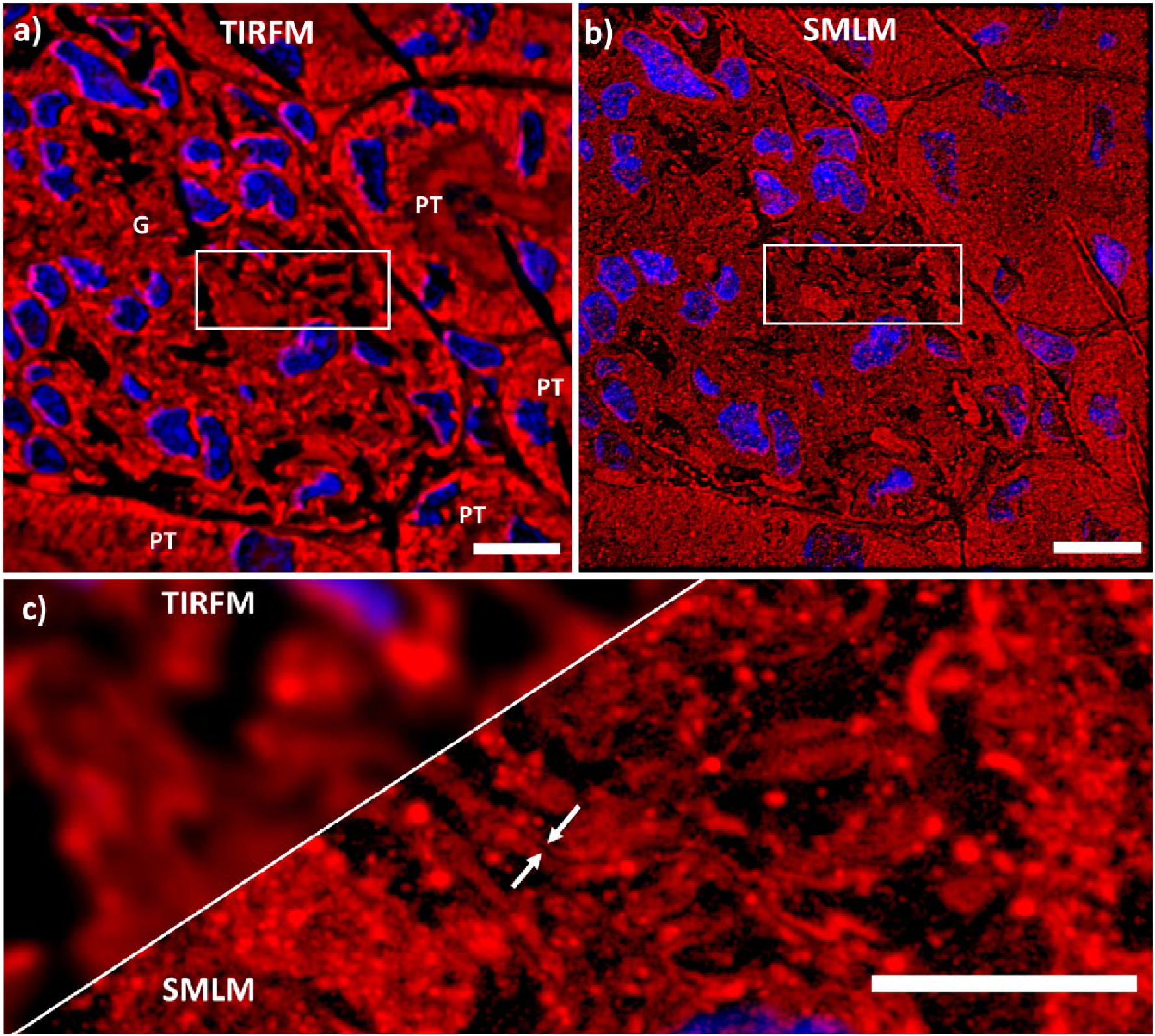
Chip-based single-molecule localization microscopy of a 400 nm mouse kidney cryosections prepared by Tokuyasu method. Membranes labeled with CellMask Deep Red (pseudo-colored in red), and nuclei labeled with Sytox Green (pseudo-colored in blue). Images were collected using a 60X/1.2 NA water immersion microscope objective. **(a)** Chip- TIRFM image of a glomerulus (G) surrounded by proximal tubuli (PT). **(b)** Chip-based SMLM image reconstructed with *d*STORM algorithm. **(c)** A magnified view of the white rectangles in (a-b) allows a comparison between these two imaging techniques. In particular, the white arrows in the SMLM segment show two lines with an average separation of 100 nm that is otherwise not observable in the TIRFM segment. Arguably, this ultrastructural feature is in agreement with the dimensions of the glomerular basal membrane present in this filtration compartment of the kidney. Scale bars (a-b) 10 μm, (c) 5 μm.

Chip-based SMLM/*d*STORM supports three to four-fold resolution improvement over diffraction limited imaging using a standard upright optical microscopy set-up with a slight modification. Moreover, the chip-based SMLM/*d*STORM approach benefits from the inherent advantage of decoupled illumination and collection light paths, which allows a user-defined choice of imaging objective lens without altering the TIRF excitation delivered by the chip. With further efforts in immunolabeling (see *Supplementary Information S9*) and system automation, chip-based SMLM could dramatically shorten the diagnostic time of nephrotic diseases that, up to now, are identified via low-throughput and expensive methods such as electron microscopy. While chip-based illumination enables the imaging of large areas, the essential challenge of SMLM relies on the need for a large number of frames for the reconstruction of a super-resolved image. Therefore, for routine histopathological analysis, it is opportune to explore alternative imaging methods, e.g. IFON, with lower demands in the number of frames necessary for super-resolution.

### 4.3 Chip-based IFON imaging

To achieve a shorter acquisition time while maintaining imaging of large areas with improved contrast and resolution, we explored chip-based intensity fluctuation optical nanoscopy (IFON) of tissue samples. IFON comprises a set of techniques that exploit the photokinetic properties of fluorescent molecules to resolve structures beyond the diffraction limit of optical microscopes [39]. The techniques examine the stochastic emission of fluorophores through statistical analysis of the intensity levels of a given image stack, allowing the identification of fluorescent emitters with sub-pixel precision. Among the IFON techniques, the multiple signal classification algorithm (MUSICAL) [14] stands out as a promising tool for fast and reliable image reconstruction of biological data [39], achieving sub-diffraction resolution through low excitation intensities, fast acquisition, and relatively small datasets (100 – 1000 frames per image stack).

The main challenges to implement IFON on histological samples are the high density and heterogeneity of the tissue samples. The spatio-temporal fluctuations are a decreasing function of the spatial density of the labels. In other words, a high density of labels results in a higher average signal at the cost of low variance in the fluorescence intensity over time. As a consequence, typically the IFON techniques are demonstrated on fine sub-cellular structures (e.g. actin filaments, microtubules, and mitochondria) fluorescently labeled on plated cells. Thus, densely labeled structures such as endoplasmic reticulum or lipid membranes are generally avoided. Tissue samples, with a higher density of labels, put even stronger demands on computational algorithms. Here, instead of relying only on the intrinsic fluctuations of the fluorophores, we propose to exploit also the intensity variations induced by the multi-mode interference (MMI) pattern (speckle-like illumination) generated by the photonic chip (see *Supplementary Information S4*). In this approach, on-chip MMI illumination patterns are modulated over time by scanning the illumination spot over the waveguide input facet. This modulates the fluorescence emissions from the fluorophores with the spatial intensity distribution of the illumination pattern at any given time. Due to the constructive and destructive interferences, bright and dark regions are formed, artificially introducing sparsity in the spatiotemporal fluctuations. In addition, due to the high refractive index of the waveguide core (*n* = 2.1 for Ta_2_O_5_ and *n* = 2 for Si_3_N_4_), the MMI pattern obtained on top of the waveguide surface are sub-diffraction limit and thus carry higher spatial frequencies than what can be obtained using free-space optics [15]. Here, we used such on-chip engineered illumination for super-resolution imaging using the MUSICAL method.

To interrogate the capability of the photonic chip for IFON-based imaging of histological samples, we used chorionic villi tissue cryosections from human placenta. For IFON studies, we focused on the visualization of ultrastructural features in the microvilli. The microvilli are actin-based membrane protrusions that increase the contact area between the syncytiotrophoblastic cells and the maternal blood, facilitating the biochemical exchange between the maternal and the fetal side, and supporting mechano-sensorial functions of the placenta [40]. Due to the physical dimensions of these structures (on average, 100 nm in diameter and 500 nm in length [41, 42]), and their tight confinement along the apical side of the syncytiotrophoblastic cells, the morphological features of the microvilli are not discernible through conventional optical microscopy and, therefore, represent an ideal element to benchmark the resolution possibilities offered by chip-based IFON.

The samples were prepared and imaged with a 60X/1.2NA microscope objective following the steps described for *Chip-based multicolor TIRFM imaging*. To avoid unspecific background signal, only the F-actin and nuclei markers were used (Phalloidin-Atto565 and Sytox Green, respectively). Further, the 500-frames image stack corresponding to the F-actin was analyzed with MUSICAL, resulting in a super-resolved and improved contrast image over a field of view of 220 × 220 μm^2^ (Figure 4a). The implementation of a soft thresholding scheme in MUSICAL [43] allowed the identification of individual microvilli along the syncytiotrophoblast's brush border (Figure 4d), which were otherwise unclear in the averaged chip-TIRFM image (Figure 4b). The resolution enhancement of MUSICAL is quantified through line-profile measurements over two adjacent microvilli. Where chip-TIRFM image (Figure 4c) showed two structures merged as a single element, the MUSICAL reconstruction (Figure 4e) revealed the separation between them. On-chip MUSICAL not only increases the resolution but improves the contrast of the image, which is a valuable parameter during visual investigations by histopathologists.

**Figure 4.**
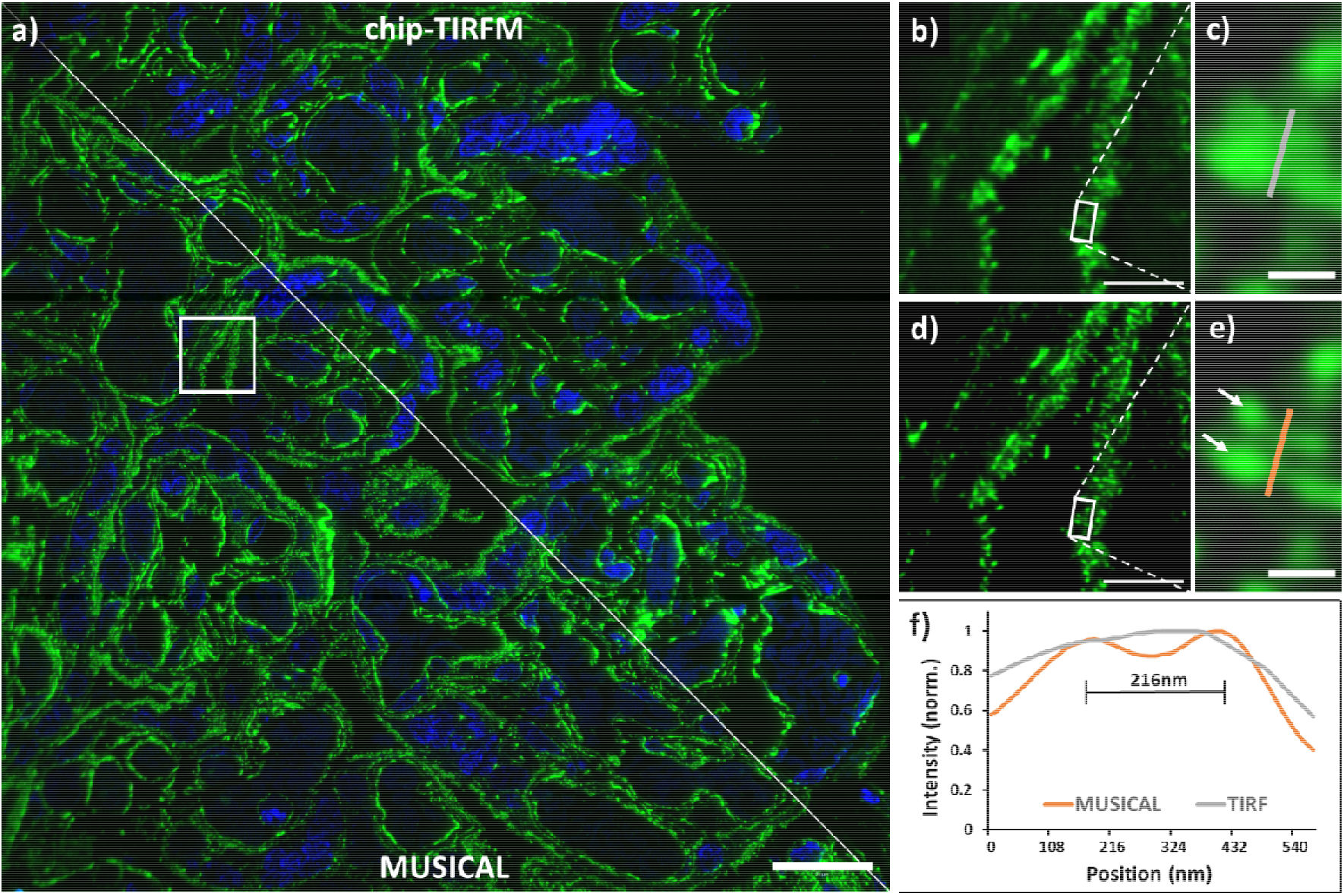
Chip-based intensity fluctuation optical nanoscopy of a 400 nm thick placental tissue section prepared per Tokuyasu method. F-actin labeled with Phalloidin-Atto565 (pseudo-colored in green), and nuclei labeled with Sytox Green (pseudo-colored in blue). **(a)** Multicolor fluorescent image over a 220 × 220 μm^2^ FOV acquired with a 60x/1.2NA microscope objective. A solid white line divides the image into two segments, illustrating the averaged chip-TIRFM on the top and the MUSICAL reconstruction at the bottom. **(b,d)** A magnified view of the white box in (a) allows for visualization of the microvilli (MV) lining the syncytiotrophoblast's brush border. **(c)** Further magnification of the white box in (b) shows a single structure. **(e)** White arrows denote the location of two adjacent MV over the magnified white box in (d). **(f)** Line-profile measurements reveal a separation of 216 nm between two adjacent MV on the MUSICAL reconstruction in (e) that is otherwise not distinguishable on the averaged chip-TIRFM image in (c). Scale bars (a) 20 μm, (b,d) 5 μm, and (c,e) 500 nm.

A recent study reported the visualization of individual microvilli with a 2-fold resolution improvement employing 3D-SIM [44]. Although several experts have proposed SIM as the fastest SRM technique for histopathology [1, 11, 45, 46], the typical FOV of this method with high magnification microscope objectives (for example, 60X/1.42NA) is about 40 × 40 μm^2^. Therefore, to match the same field of view achieved with the photonic chip, a tile mosaic of 7 × 7 SIM images would be required (see *Supplementary Information S10*). For conventional 3D-SIM, this not only implies a prolonged time for the data acquisition, but also a lengthy image reconstruction that rounds up to 2.5 h. On contrary, the MUSICAL implementation we used here was able to obtain a high-resolution image over a large field of view within a combined collection and processing time of 10 min for the 500-frames acquired on the photonic chip. From a practical perspective, the high-resolution visualization over large areas supported by chip-based IFON opens the door for improved assessment of placental microstructure both for basic research as well as for clinical assessment of placental pathologies associated with morphological changes in the microvilli, as documented in placental dysfunction disorders, such as pre-eclampsia [41].

### 4.4 Chip-based CLEM imaging

Combining the specificity of fluorescence microscopy with the high resolution of electron microscopy allows the visualization of proteins of interest along with the ultrastructural context of the tissues. Although recent reports have proposed silicon wafers for correlative light and electron microscopy (CLEM) [47, 48], they employed EPI-illumination through high-magnification microscope objectives, providing a limited field of view of the fluorescent signal. Here, we employed zebrafish eye retina cryosections of 110 nm thickness to demonstrate the compatibility of the chip platform with CLEM studies. Zebrafish is a well-established model for the study of retinal diseases [49]. The samples were prepared in the same manner as the placental and renal sections, except for the initial washing steps of the cryoprotectant. We found that the optimal washing temperature of the sucrose-methyl cellulose solution for these samples was 0°C, over an incubation time of 20 minutes (see detailed protocol in Materials and Methods). Three structures were labeled for the study: a) the F-actin filaments, b) the nuclei, and c) the outer mitochondrial membrane. The first two structures were labeled through direct markers using Texas Red-X Phalloidin and Sytox Green, respectively, whereas the latter was labeled by immunofluorescence using rabbit anti-Tomm20 as a primary antibody, and Alexa Fluor 647-conjugated donkey anti-rabbit as a secondary antibody.

The samples were first imaged in chip-based TIRFM mode for each channel using a 60X/1.2NA to obtain a diffraction-limited multicolor image (**Figure 5a**). Thereafter, the sections were platinum-coated and imaged on a scanning electron microscope over the same region of interest (**Figure 5b**). A magnified view of the TIRFM image (**Figure 5c**) allows for the observation of the F-actin filaments lining the outer segments of the photoreceptors (in green), as well as the mitochondria clusters (in magenta), and the location of the nuclei (in cyan). The same TIRFM dataset is used for post-processing through MUSICAL, allows for a precise correlation of both the F-actin and the mitochondrial signals with the corresponding SEM image (**Figure 5d,e,f**).

**Figure 5.**
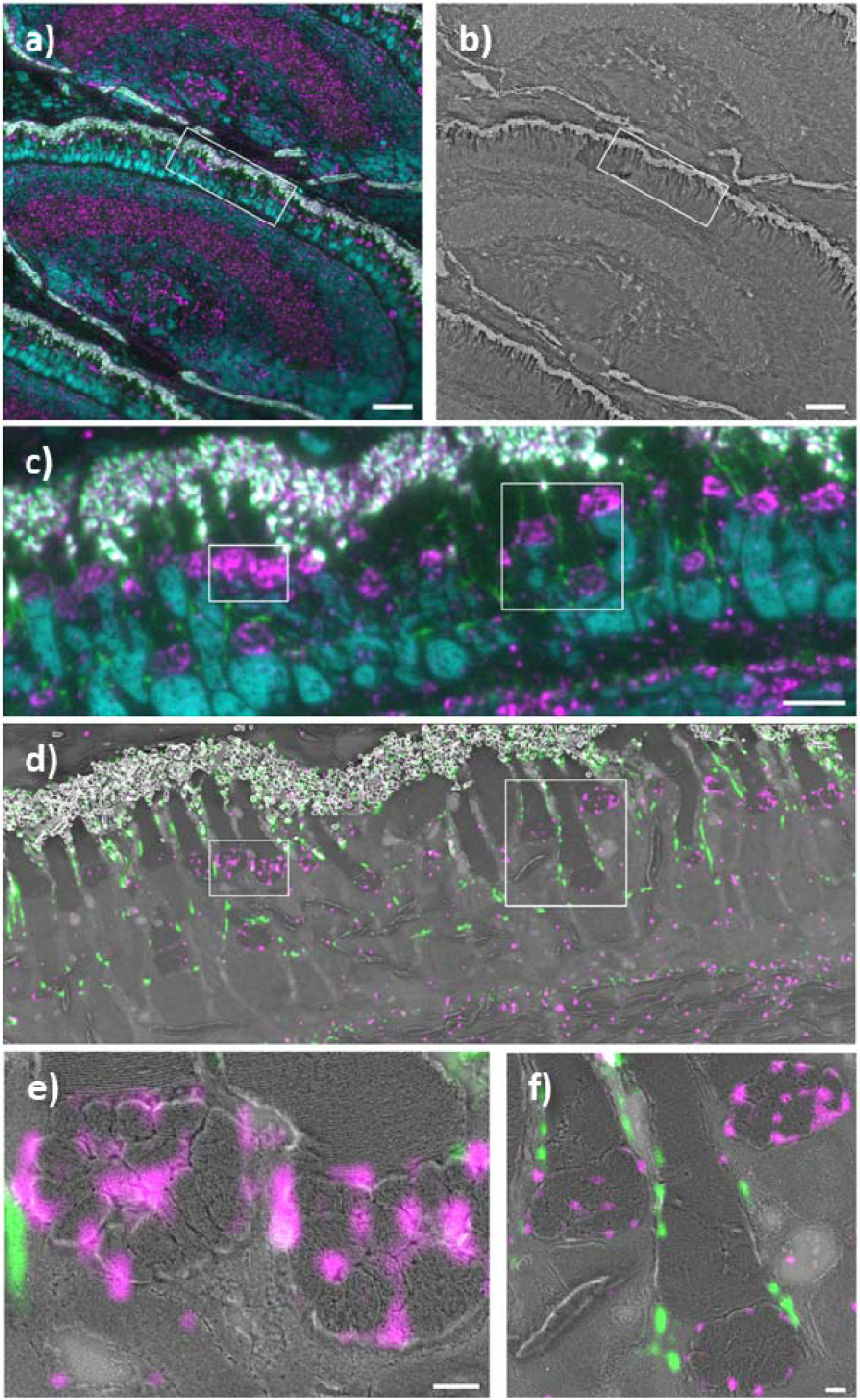
Chip-based CLEM imaging of a 110 nm thick zebrafish retina cryosections prepared by Tokuyasu method on a 600 μm wide optical waveguide. **(a)** Diffraction-limited chip-TIRFM image. In magenta, mitochondrial clusters immunolabeled with rabbit anti Tomm20 protein (primary antibody) and Alexa Fluor 647-conjugated donkey anti-rabbit (secondary antibody). In green, actin segments labeled with Texas Red-X Phalloidin. In cyan, nuclei labeled with Sytox Green. **(b)** Scanning electron microscope image of the same region shown in (a) scanned at 30 nm pixel size. **(c)** high magnification image of the white frame in (a) showing the diffraction-limited chip-TIRFM signal of mitochondria, actin, and nuclei. **(d)** CLEM image of areas in frames (a) and (b). Scanning electron microscope image acquired at 4 nm pixel size correlates with the MUSICAL images of mitochondria (magenta) and actin (green). **(e)** CLEM image of the white region in (d). MUSICAL image of the Tomm20 signal (magenta) in the outer membrane of mitochondria correlating with the morphology of the complex clusters of mitochondria. The tightly packed membranes of the outer segment are clearly recognized. **(f)** CLEM image of MUSICAL-processed actin signal along with the outer segments (green) and three mitochondria clusters. The MUSICAL signal in (d), (e) and (f) were gamma-corrected to increase the contrast of the actin (γ=1.2) and the Tomm20 signal (γ=1.1). Scale bars (a,b) 20 μm, (c,d) 5 μm, (e,f) 500 nm.

Notably, the waveguide widths on the chip not only accommodated the whole zebrafish retina but also allowed the observation of several serial sections in a ribbon (see *Supplementary Information S12*). Also, the combination of the thin section thickness of the Tokuyasu samples with the limited extent of the evanescent field dramatically improved the axial resolution of the fluorescent signal, enabling high-contrast images. Put together, these features are advantageous for confirming signal specificity throughout different subcellular compartments, opening up the possibility for 3D-stacking via serial section imaging [50, 51]. Moreover, the flatness of the chip serves as an optimal platform for SEM, allowing autonomous imaging over large areas. A simple thin layer of platinum deposited on top of the chip minimizes the charging effects and enables a good correlation between the light and the electron microscopy images. Importantly, the photonic chip can incorporate coordinate land-markings to facilitate the location and further correlation of the ROIs under study [29]. Lastly, the chip-based CLEM strategy presented here, in combination with the Tokuyasu method, can be executed within one working day from the sample sectioning steps to the SEM imaging, implying a significant time improvement as compared to the typical one-week imaging throughput associated with most CLEM approaches [34].

## 5. Conclusions and future perspectives

In this study, we demonstrated the capabilities of the photonic chip as a feasible imaging platform for morphological assessment of thin Tokuyasu sections of a variety of tissues. The photonic chip-based microscopy technique offers several advantages for histopathology: a) it allows a broad range of imaging modalities over large fields of view including TIRFM, SMLM, IFON, and CLEM using a single standard optical microscope set-up; b) the imaging process can be seamlessly performed on conventional optical microscopes upon some modifications; and c) the photonic chip withstands all the chemical incubations and thermal conditions associated with the sample preparation. These features make the photonic chip an attractive platform for fluorescence-based histopathological studies where high-throughput, high-contrast, and high-resolution are crucial for the diagnosis of diseases. We anticipate that, upon specific labeling and image processing efforts, the photonic chip could assist both in reducing the processing time and in improving the assessment quality of pathologies that –to date– require ultrastructural observation using electron microscopy. Additionally, in CLEM experiments, the photonic chip could be used for fast assessment of ultrastructural preservation in tissues.

The photonic chip approach also reduces the complexity of the optical nanoscopy setups by miniaturization of the excitation light path, simplifying the implementation of multimodal imaging and facilitating a larger adoption of super-resolution microscopy in clinics and hospitals. In addition, the photonic chip can be mass-produced through standard semi-conductor lithography processes, benefitting from low-cost manufacturing scalability. We foresee that further developments in coupling automation and the integration of microfluidics systems could dramatically improve the performance of the photonic chip platform, enabling more efficient and repeatable labeling, as well as fast multiplexed imaging. Moreover, the implementation of advanced labeling strategies such as DNA-PAINT [27] and Exchange-PAINT [52] can effectively reduce the background signal, improving resolution, and support multiplexed acquisition. Also, on-chip technology facilitates the integration of other on-chip optical functions such as Raman spectroscopy [53], waveguide trapping [54], microfluidics [55], phase microscopy [24], among others.

While the photonic chip illumination strategy allows excitation over large areas, e.g. several centimeters in the present case, the light collection area is presently limited by the collection objective lens. Thus, it can be envisioned that the integration of microlens arrays [56] for light collection will open avenues that would make on-chip technology capable of handling the high-throughput imaging needed for routine histopathology. Moreover, the photonic chip can be designed and manufactured into standard microscope glass slide dimensions, allowing for a fully automated sample preparation through commercially available immunoassay analyzers, or via novel microfluidic techniques for multiplex immunofluorescence staining of clinically relevant biomarkers [57].

Despite the encouraging imaging results obtained in this study, we acknowledge that the Tokuyasu samples represent a minority among the available histological methods. Also, we are aware that the maximum section area possible with the Tokuyasu cryosections (500 × 500 μm^2^) may be insufficient for large-scale histopathological evaluation. However, this is an inherent limitation imposed by the sample preparation technique rather than the photonic chip imaging surface. Future chip-based histology studies should address the compatibility of this microscopy platform with widely accessible samples including formalin-fixed paraffin-embedded (FFPE) and cryostat-sliced sections.

## 6. Materials and Methods

### 6.1 Photonic chip description and fabrication

The photonic chip is composed of three layers: i) a bottom silicon (Si) substrate, ii) an intermediate cladding of silicon dioxide (SiO_2_), and iii) a top waveguide layer of a high refractive index material made of either silicon nitride (Si_3_N_4_, *n* = 2.0) or tantalum pentoxide (Ta_2_O_5_, *n* = 2.1) (see Figure 1a). The high refractive index contrast (HIC) between the waveguide materials and the adjacent imaging medium and sample (*n* ≈ 1.4), allows the confinement and propagation of the excitation light via total internal reflection (TIR), enabling chip-based total internal reflection fluorescence microscopy (chip-TIRFM) (Figure 1c). Diverse geometries have been previously studied for chip-TIRFM, including slab, rib, and strip waveguides [15]. Here, we chose uncladded strip waveguides with heights ranging from 150 nm to 250 nm and widths varying from 200 μm to 1000 μm (see Figure 1b).

In this study, we used both Si_3_N_4_ and Ta_2_O_5_ chips for chip-TIRFM imaging of tissue sections. These were fabricated in distinct places: i) the Si N waveguide chips were manufactured according to CMOS fabrication process at the Institute of Microelectronics Barcelona (IMB-CNM, Barcelona, Spain) as detailed elsewhere [15, 17, 58]; ii) the Ta_2_O_5_ chips were manufactured at the Optoelectronics Research Center (ORC, University of Southampton, UK), following the process herewith detailed [59]. Waveguides of 250 nm thickness were fabricated by deposition of Ta_2_O_5_ film on a commercially-available 4” Si substrate having a 2.5 μm thick SiO_2_ lower cladding layer (Si-Mat Silicon Materials, Germany) using a magnetron sputtering system (Plasmalab System 400, Oxford Instruments). The base pressure of the Ta_2_O_5_ deposition chamber was kept below 1 × 10^−6^ Torr with Ar:O_2_ flow rates of 20 sccm : 5 sccm and the substrate temperature was maintained at 200 °C throughout the deposition process. Photolithography was used to create a photoresist mask for further dry etching to fabricate strip waveguides. First, 1 μm thick positive resist (Shipley, S1813) was coated on top of a 250 nm Ta_2_O_5_ film and then prebaked (1 × 30 min) at 90 °C. Then, the wafer was placed into a mask aligner (MA6, Süss MicroTec), and illuminated with the waveguide pattern. The Ta_2_O_5_ layer, which was not covered with photoresist, was fully etched to obtain strip waveguides of 250 nm height using an ion beam system (Ionfab 300+, Oxford Instruments) fed with argon at a flow rate of 6 SSCM. The process pressure (2.3×10^−4^ Torr), beam voltage (500 V), beam current (100 mA), radiofrequency power (500 W), and substrate temperature (15 °C) were kept constant. Finally, the wafers were placed in a 3-zone semiconductor furnace at 600 °C in an oxygen environment for 3 hours (in batch) to reduce the stress and supplement the oxygen deficiency created in Ta_2_O_5_ during the sputtering and the etching process [23].

Upon reception, the wafers were split into individual chips using a cleaving system (Latticegear, LatticeAx 225). The remaining photoresist layer from the manufacturing process was removed by immersion in acetone (1 × 1 min). The chips were then cleaned in 1% Hellmanex in deionized water on a 70 °C hotplate (1 × 10 min), followed by rinsing steps with isopropanol and deionized water. The chips were finally dried with nitrogen using an air blowgun. To improve the adhesion of the tissue sections, the chips were rinsed with 0.1 % w/v poly-L-lysine solution in H_2_O and let dry in a vertical position (1 × 30 min).

### 6.2 Sample collection and preparation

#### 6.2.1 Ethical statement

Both animal and human samples were handled according to relevant ethical guidelines. Healthy placental tissues were collected after delivery at the University Hospital of North Norway. Written consent was obtained from the participants following the protocol approved by the Regional Committee for Medical and Health Research Ethics of North Norway (REK Nord reference no. 2010/2058-4). Treatment and care of mice and pigs were conducted following the guidelines of the Norwegian Ethical and Welfare Board for Animal Research. Zebrafish experiments were conducted according to Swiss Laws and approved by the veterinary administration of the Canton of Zurich, Switzerland.

#### 6.2.2 Preparation of Tokuyasu sections for chip-based TIRFM, IFON, and SMLM

Human placental and murine (NZBxNZW)F1 kidney tissue samples were cryopreserved following the Tokuyasu method for ultracryotomy described elsewhere [44, 60]. In short, biopsies blocks of approximately 1 mm^3^ were collected, rinsed in 9 mg/mL sodium chloride, fixed in 8% formaldehyde at 4°C overnight, infiltrated with 2.3M sucrose at 4°C overnight, mounted onto specimen pins, and frozen in liquid nitrogen. Thereafter, the samples were transferred to a cryo-ultramicrotome (EMUC6, Leica Microsystems) and sectioned with a diamond knife into thin slices ranging from 100 nm to 1 μm thickness. The sections were collected with a wire loop containing a 1:1 cryoprotectant mixture of 2% methylcellulose and 2.3 M sucrose and transferred to photonic chips coated with poly-L-lysine and equipped with custom-made polydimethylsiloxane (PDMS) chambers of approximately 130 μm-height [17] (Figure 1b). The samples were stored on Petri dishes at 4°C before subsequent steps.

Diverse staining strategies were employed according to each imaging modality:

i. For *Chip-based multicolor TIRFM* imaging, human placental sections of 400 nm were direct-labeled for membranes, F-actin, and nuclei as described herewith. First, the cryoprotectant mixture was dissolved by incubating the samples in phosphate-buffered saline (PBS) (3 × 10 min) at 37 °C. Thereafter, the samples were incubated in a 1:2000 solution of CellMask Deep Red in PBS (1 × 15 min) at room temperature (RT) and subsequently washed with PBS (2 × 5 min). Next, the sections were incubated in 1:100 Phalloidin-Atto565 in PBS (1 × 15 min) and washed with PBS (2 × 5 min). Further, the samples were incubated in 1:500 Sytox Green in PBS (1 × 10 min) and washed with PBS (2 × 5 min). Finally, the sections were mounted with #1.5 coverslips using Prolong Diamond and sealed with Picodent Twinsil.
ii. For *Chip-based SMLM imaging*, mouse kidney cryosections of 400 nm were labeled for membranes and nuclei using CellMask Deep Red and Sytox Green, respectively, following identical concentrations and incubation steps as for the *Chip-based multicolor TIRFM imaging* experiments. To enable photoswitching of the fluorescent molecules, the samples were mounted with a water-based enzymatic oxygen scavenging system buffer as described in previous works [15, 16]. Thereafter, the sections were covered with #1.5 coverslips and sealed with Picodent Twinsil.
iii. For *Chip-based IFON imaging*, human placental sections of 400 nm were prepared identically to the *Chip-based multicolor TIRFM imaging* experiment, except for the membrane labeling and subsequent washing steps that were omitted. In all cases, the labeled cryosections were stored at 4°C and protected from light before imaging. *Supplementary Information S8* provides a detailed description of the materials and reagents used in this protocol.

#### 6.2.3 Preparation of Tokuyasu sections for chip-based CLEM

For Chip-based CLEM imaging, zebrafish eyes were prepared as described elsewhere [61]. Briefly, 5 days-post-fertilization larvae were euthanized in tricaine and fixed with 4 % formaldehyde and 0.025% glutaraldehyde in 0.1 M sodium cacodylate buffer (1 × 16 h) at 4 °C. Subsequently, eyes were dissected and washed in PBS, placed in 12 % gelatin (1 × 10 min) at 40 °C, and finally left to harden at 4°C. Embedded eyes were immersed in 2.3 M sucrose and stored at 4°C before further storage in liquid nitrogen. Ultrathin sections of 110 nm thickness were obtained with a cryo-ultramicrotome (Ultracut EM FC6, Leica Microsystems) using a cryo-immuno diamond knife (35° - size 2 mm, Diatome). The cryosections were transferred to photonic chips fitted with a PDMS frame and stored at 4 °C before staining. The samples were incubated in PBS (1 × 20 min) at 0 °C, followed by two washing steps in PBS (2 × 2 min) at RT to dissolve the cryoprotectant. Then, the samples were pre-incubated with a blocking solution (PBG) for 5 min, followed by incubation (1 × 45 min) in a 1:50 solution of rabbit anti-Tomm20 in PBG blocking buffer at RT. After several rinsing (6 × 2 sec) and washing (1 × 5 min) in PBG, the specimens were incubated (1 × 45 min) with an Alexa Fluor 647-conjugated secondary donkey anti-rabbit antibody at 1:200 concentration in PBG at RT. For the acting staining, the samples were washed in PBS (6 × 1 min), followed by incubation with Texas Red-X Phalloidin (1 × 10 min) at 1:50 concentration in PBS. After washes in PBS (2 × 5 min), the samples were incubated in a 1:500 solution of Sytox Green nuclear staining in PBS (1 × 10 min), followed by washes in PBS (2 × 5 min), and mounting with a 1:1 mixture of PBS and glycerol (49782, Sigma-Aldrich) and covered with a #1.5 glass coverslip before chip-TIRFM imaging. *Supplementary Information S8* provides a detailed description of the materials and reagents used in this protocol.

### 6.3 Chip imaging and processing

#### 6.3.1 Chip-based imaging

The chip-TIRFM setup was assembled using a modular upright microscope (BXFM, Olympus), together with a custom-built photonic chip module as shown in Figure 1c and *Supplementary Information S11*. A fiber-coupled multi-wavelength laser light source (iChrome CLE, Toptica) was expanded and collimated through an optical fiber collimator (F280APC-A, Thorlabs) to fill the back aperture of the coupling MO (NPlan 50X/NA0.5, Olympus). Typical illumination wavelengths used were λ_1_ = 640 nm, λ_2_ = 561 nm, and λ_3_ = 488 nm. Both the optical fiber collimator and the coupling objective were mounted on an XYZ translation stage (Nanomax300, Thorlabs) fitted with an XY piezo-controllable platform (Q-522 Q-motion, PI) for fine adjustments of the coupling light into the waveguides. The photonic chips were placed on a custom-made vacuum chuck fitted on an X-axis translation stage (XRN25P, Thorlabs) for large-range scanning of parallel waveguides. Fluorescent emission of the samples was achieved via evanescent field excitation upon coupling of the laser onto a chosen waveguide, as detailed elsewhere [15] (Figure 1a,c). Various MO lenses were used to collect the fluorescent signal, depending on the desired FOV, magnification, and resolution (4X/0.1NA, 20X/0.45NA, and 60X/1.2NA water immersion). An emission filter set composed of a long-pass filter and a band-pass filter was used to block out the excitation signal at each wavelength channel (see *Supplementary Information S11* for details). The emission signal passed through the microscope’s 1X tube lens (U-TV1X-2, Olympus) before reaching the sCMOS camera image plane (Orca-flash4.0, Hamamatsu). Both the camera exposure time and the laser intensity were adjusted according to the experimental goal. For TIRFM imaging, the camera exposure time was set between 50 ms and 100 ms, and the input power was incrementally adjusted until the mean histogram values surpassed 500 counts. For SMLM, the acquisition time was set to 30 ms while the input power was set to its maximum level to enable photoswitching. Depending on the coupling efficiency, typical input powers were between 10% and 60% for TIRFM imaging, and between 90% to 100% for SMLM imaging. To reduce photobleaching of the fluorescent markers, the image acquisition was sequentially performed from less energetic to more energetic excitation wavelengths. To deal with the anisotropic mode distribution of the multi-mode interference pattern at the waveguide, the coupling objective was laterally scanned at < 1 μm steps over a 50 μm – 200 μm travel span along the input facet of the chip while individual images were taken. Image stacks of various sizes were acquired according to the imaging technique. Typically, 100 – 1000 frames for TIRFM and 30000 – 50000 frames for SMLM. White light from a halogen lamp (KL1600 LED, Olympus) was used for bright-field illumination to identify the regions of interest (ROI) through the collection objective. To reduce mechanical instability, the collection path of the system was fixed to the optical table, while the photonic chip module was placed onto a motorized stage (8MTF, Standa) for scanning across the XY directions. An optical table (CleanTop, TMC) was used as the main platform for the chip-TIRFM setup. *Supplementary Information S11* offers a detailed description of the chip-TIRFM setup.

#### 6.3.2 CLEM imaging

After chip-TIRFM imaging, both the coverslip and the PDMS frame were removed and the samples fixed with 0.1% glutaraldehyde. Thereafter, the samples were incubated with methylcellulose followed by centrifugation at 4700 rpm (Heraeus Megafuge 40R, Thermo Scientific) in a falcon tube. After drying (2 × 10 min) at 40 °C on a heating plate, the photonic chips were transferred to an electron beam evaporator (MED 020, Leica Microsystems). The specimen was then coated with platinum/carbon (Pt/C, 10⍰nm) by rotary shadowing at an angle of 8⍰degrees [48]. Thereafter, the photonic chips were mounted on a 25 mm Pin Mount SEMclip (#16144-9-30, Ted Pella) and imaged at 4 nm pixel size with a scanning electron microscope (Auriga 40 CrossBeam, Carl Zeiss Microscopy) at a low-accelerating voltage (1.5 keV). *Supplementary Information S12* illustrates various steps of SEM imaging on a photonic chip.

#### 6.3.3 Image processing

The acquired frames were computationally processed on the open-source software Fiji [62] according to the desired imaging technique. To obtain diffraction-limited TIRFM images, the image stacks were computationally averaged using the *Z Project tool*. Thereafter, the averaged images were deconvolved with the *DeconvolutionLab2* plugin [63], using a synthetic 2D point spread function (PSF) matching the effective pixel size of the optical system. Lastly, the *Merge Channels* tool was used to merge and pseudocolor independent averaged channels into a multicolor composite TIRFM image. SMLM images were reconstructed using the thunderSTORM plugin [64]. For CLEM, the acquired TIRFM stacks were first processed with the NanoJ SRRF plugin [65] and then correlated with the EM images using the TrakEM2 plugin [66].

## 7. Acknowledgments and funding

The authors thank the collaborators at UiT The Arctic University of Norway, including Randi Olsen for providing the cryosections, and Deanna Wolfson for her valuable labeling recommendations. The authors also acknowledge Åsa Birna Birgisdottir and Trine Kalstad, for providing the pig heart samples, and Prof. Dr. Stephan Neuhauss, University of Zurich, for providing the zebrafish eye samples. The authors would like to express their appreciation to Prof. James Wilkinson (University of Southampton) and Dr. Senthil Murugan Ganapathy (University of Southampton) for discussions on the waveguide platform fabrication. B.S.A. acknowledges the funding from the Research Council of Norway, project # NANO 2021–288565 and project # BIOTEK 2021–285571.

## 8. Author contribution

B.S.A. conceived the idea and supervised the project. L.E.V.H. and V.D. planned and coordinated the experiments, performed sample labeling, chip-TIRFM imaging, and post-processing of the data. J.C.T. and V.D. built the chip-based microscope setup. S.A. and K.A. performed MUSICAL reconstruction. D.C., V.D., and L.E.V.H. performed SMLM acquisition. D.C. performed the *d*STORM reconstruction. L.E.V.H., J.C.T., and J.M.M. performed chip-TIRFM imaging of the zebrafish eye. G.B. provided the zebrafish cryosections. G.B, J.M.M., and U.Z. designed the experimental conditions for SEM imaging. J.M.M. performed the SEM imaging for CLEM. F.T.D. and A.P. respectively designed and fabricated the Ta_2_O_5_ photonic chips. G.A. provided the human placental sample. M.N. collected and preserved the human placental samples. G.A. and M.N. helped with the placental image interpretations. A.K.H. and K.A.F. collected and preserved the mouse kidney tissue and assisted with the renal image interpretations. L.E.V.H. and V.D. analyzed the data, prepared the figures. L.E.V.H., V.D., and B.S.A. wrote the manuscript. All authors contributed to writing and revising selected sections of the manuscript.

## Supplementary Information

### S1 Comparison of super-resolution techniques

The relative benefits and limitations of different super-resolution techniques are summarized in Table S1. SIM: structured illumination microscopy; IFON: intensity fluctuation-based optical nanoscopy; STED: stimulated emission depletion microscopy; SMLM: single-molecule localization microscopy; EM: electron microscopy.

**Table S1.** Overview of technical capabilities of different super-resolution methods for tissue imaging Legend color: dark green = good; light green = acceptable; yellow = medium/moderate; red = problematic

|  | **SIM** | **IFON** | **STED** | **SMLM** | **EM** |
| --- | --- | --- | --- | --- | --- |
| **Resolution (nm)** | 100 | 50-150 | 20-70 | 10-55 | 1-50 |
| **Throughput (area imaged/time)** |  |  |  |  |  |
| **Multicolor Imaging** | Possible | Possible | Demanding | Demanding | N.A. |
| **Widefield/ Point scanning** | Widefield | Widefield | Point scanning | Widefield | Point Scanning |
| **Choice of Fluorophores** | Bright & photo-stable | Common | STED-capable | Photoactivable | N.A. |
| **Sample preparation** | Easy | Easy | Moderate | Moderate | Complex |
| **Cost and Complexity** | High | Low | High | Low | Very High |
| **Post-acquisition reconstruction** | Yes | Yes | No | Yes | No |

### S2 Evanescent field simulations

For the estimation of the waveguide parameters such as surface intensity and extent of evanescent field, simulations with Fimmwave (Photon Design) were performed for a strip waveguide having a width of 200 μm. Figure S2a shows the schematic diagram of a strip waveguide. The waveguides are fabricated on the SiO_2_ coated Si substrate. Figure S2b shows the distribution of the fundamental TE mode along the width and core thickness of the waveguide structure. The coupled light propagates through the length of the waveguide generating an evanescent field on its top. The surface intensity of the evanescent field depends highly on the geometry of the waveguide, refractive index differences of the core and surrounding material, and the wavelength of the coupled light. Figure S2c shows the variation in surface intensity and relative depth of the evanescent field (penetration depth) as a function of core thickness. As the core thickness increases, the surface intensity decreases dramatically. The amplitude of the penetration depth also decreases with increasing core thickness and becomes almost stable after 150 nm. The simulation results allow choosing a core thickness between 150 nm and 250 nm for chip-TIRFM applications.

**Figure S2.**
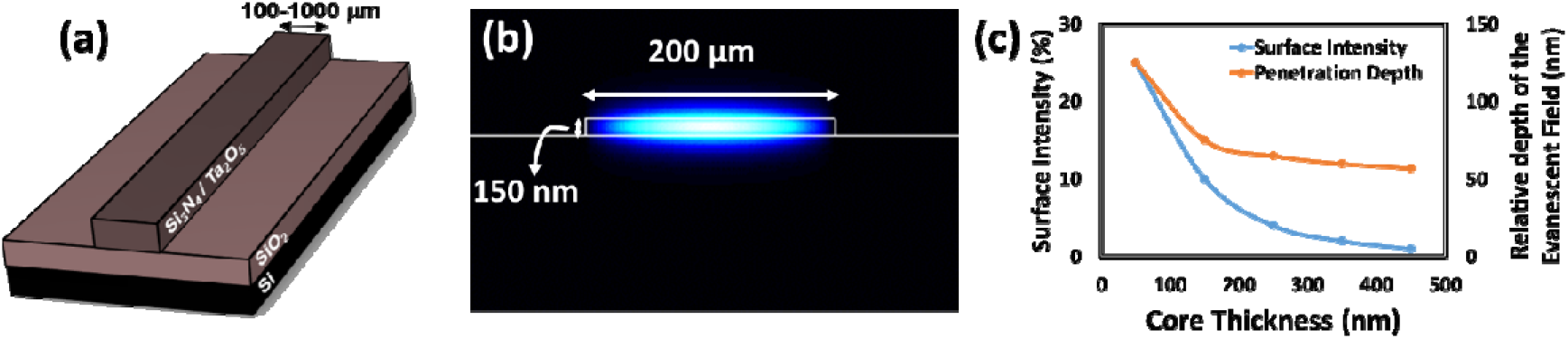
Evanescent field simulations on a Ta_2_O_5_ strip waveguide. **(a)** Schematic diagram of a photonic chip with a strip waveguide core width varying from 100 μm to 1000 μm. **(b)** Simulated field distribution of a fundamental transverse electrical (TE) mode on a Ta_2_O_5_ waveguide of 200 μm wide and 150 nm thickness. **(c)** The strength (surface intensity) and the penetration depth of the evanescent field vary as a function of the waveguide thickness. The wavelength considered in the simulation corresponds to 561 nm and the waveguide core material Ta_2_O_5_.

### S3 Sample preparation work-flow for chip-TIRFM of Tokuyasu sections

Sample preparation plays a key role in the imaging outcome of chip-based microscopy. Figure S3 provides a schematic workflow of the preparation steps for chip-TIRFM imaging of placental cryosections per the Tokuyasu method. The protocol is based on an existing procedure for fluorescent labeling of Tokuyasu sections on glass coverslip [1]. From the orange-dotted line in Figure S3a, all the steps are optimized according to the specific needs of the experiment. In particular, the washing steps of the cryoprotectant (immediately after orange box in Figure S3a) can be carried out with PBS at 37 °C instead of 4 °C PBS as detailed in Figure 2a (see more details in *Supplementary Information S5*).

**Figure S3.**
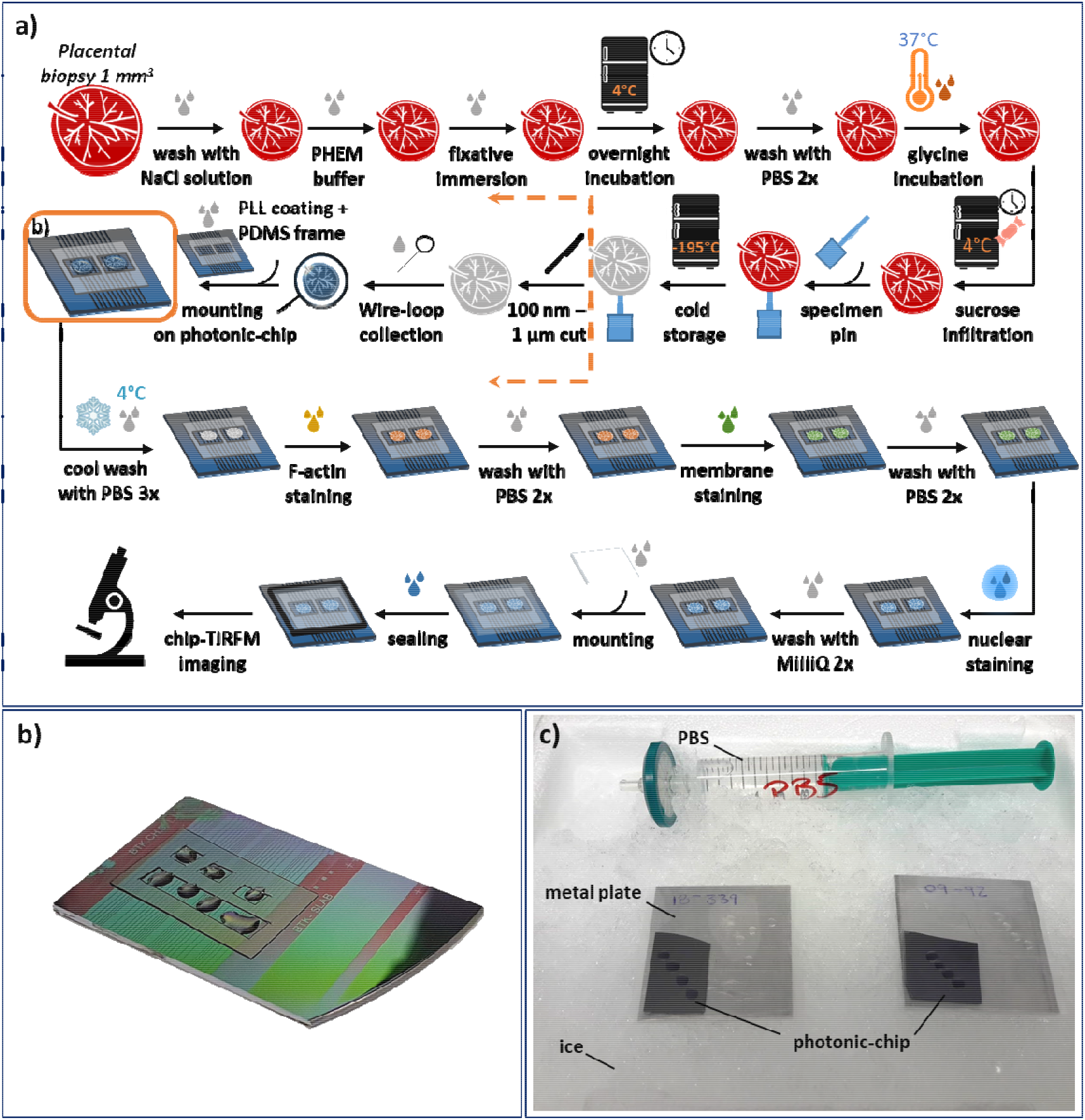
Sample preparation protocol for fluorescence staining of Tokuyasu cryosections on a photonic chip. **(a)** Schematic workflow of the sample preparation steps per the Tokuyasu method of a placental section. **(b)** Depiction of the orange box in (a) illustrating a photonic chip with Tokuyasu cryosections on top and surrounded by a custom-made PDMS frame. **(c)** The photonic chips are placed on top of metal plates on ice for the washing step in cold PBS following the orange box in (a).

### S4 Mode-averaging for homogeneous illumination in chip-TIRFM imaging

The waveguides used for tissue imaging are wide, supporting the propagation of several light modes (Figure S4a). Upon coupling of the excitation beam onto the waveguide, a non-uniform intensity distribution is observed due multiple mode interference (MMI) patterns (Figure S4b). These patterns change depending on the position of the coupling objective. To achieve isotropic illumination of the specimen, the coupling objective is scanned along the input facet of the chip while individual frames are acquired (Figure S4c). The collected image stack is averaged (Figure S4d) and then deconvolved (Figure S4e) to obtain a diffraction-limited chip-TIRFM image.

Interestingly, on-chip MMI patterns helps IFON method especially for dense tissue sample. The spatio-temporal fluctuations from the sample are a decreasing function of the spatial density of the labels. For a dense tissue sample this results in a higher average signal at the cost of low variance in the fluorescence intensity over time. This makes super-resolution imaging of tissue samples using IFON methods difficult. By using non-uniform MMI patterns not all regions (fluorophores) are excited at the same time which helps to alleviate the issues with dense labelling. Furthermore, as these MMI patterns are generated inside the photonic-chip, it carries high spatial frequencies due to high refractive index of the waveguide material.

**Figure S4.**
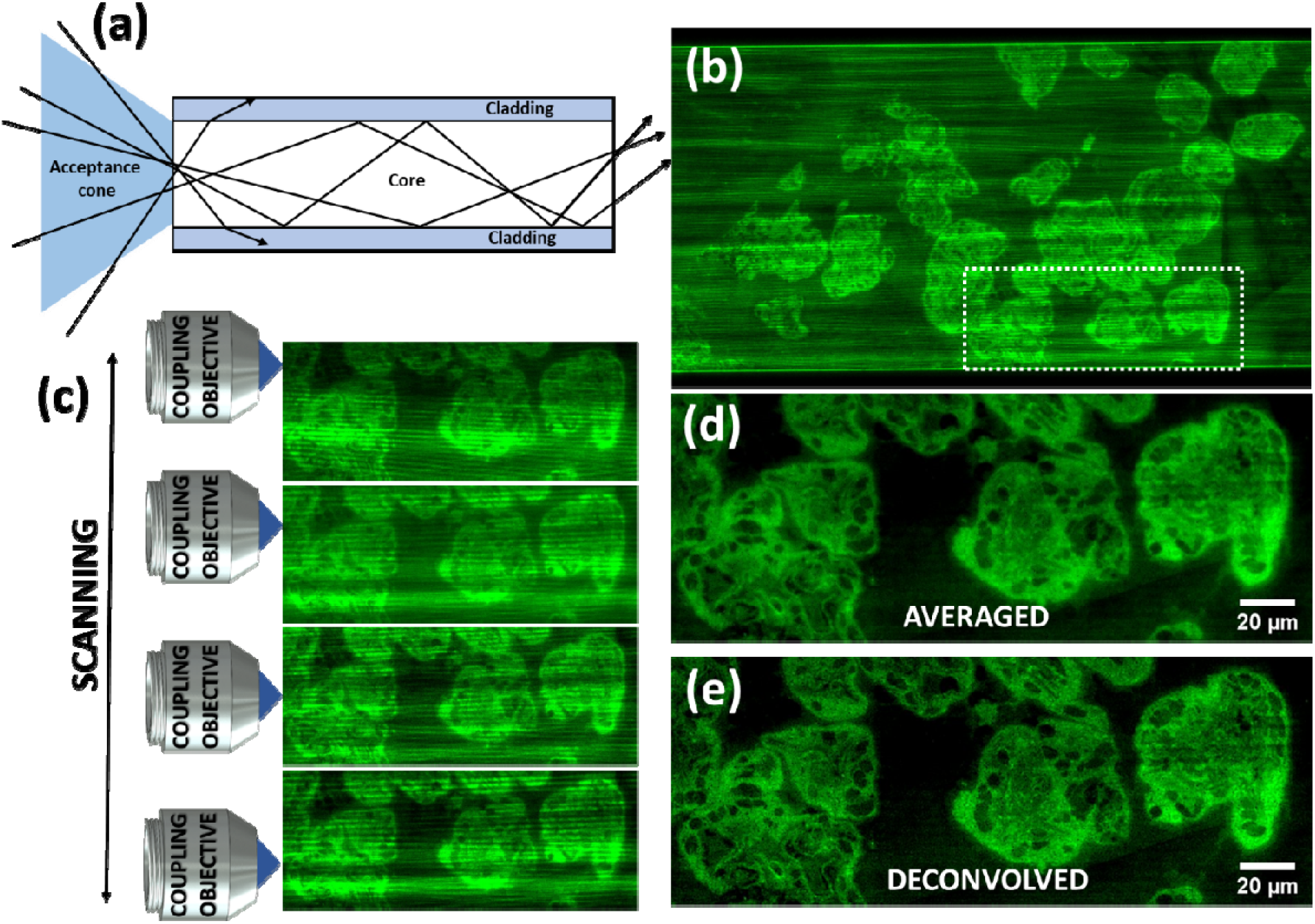
Mode-averaging for chip-TIRFM imaging of a 400 nm thick human chorionic villi cryosection. Membranes with CellMask Orange (pseudocolored in green). **(a)** Schematic diagram of a multi-mode waveguide supporting the propagation of multiple light modes. **(b)** Top view of a multi-mode pattern generated by the interference of multiple propagating light modes. **(c)** The excitation beam is scanned across the input facet of the waveguide while individual frames are collected (in this example four frames are illustrated). **(d)** The acquired stack is post-processed to obtain an averaged image with uniform intensity distribution. **(e)** The averaged image is deconvolved to obtain a high-contrast diffraction-limit chip-TIRFM image of the tissue section.

### S5 Sample preparation and imaging optimization

Despite the encouraging imaging results obtained on the photonic chip, there were experimental challenges along with this study, mainly related to sample labeling and image acquisition. In the first place, the membrane tags investigated for labeling, namely CellMask Deep Red (CMDR), CellMask Orange (CMO), DiI, and DiO, showed high affinity to the photonic chip surface, inducing a high-background signal upon coupling of the excitation light onto the waveguides. Also, the combination of F-actin markers (Phalloidin-Atto647N, Phalloidin-Atto565) together with the CellMask family dyes (CMO and CMDR) led to the appearance of labeling artifacts in the form of dense spots (see white arrows in the main manuscript, Figure 2a), hampering the quality of the acquired images. We observed that, after an initial bleaching step of the membrane dye (see *Supplementary Information S6*) the remaining fluorescent signal at the sample was generally stable through the image acquisition, allowing continuous imaging over prolonged timescales (> 5 min). We hypothesize that due to the exponential decay of the evanescent field (see *Supplementary Information S2*), the fluorophores in close vicinity to the chip are more susceptible to irreversible photodamage than those located further away, hence allowing for localized bleaching at the waveguide-sample interface. Remarkably, no labeling artifacts were observed when F-actin markers were used separately from their membrane counterparts (see Figure 4 and Figure 5 in the main manuscript).

Next, the decaying nature of the evanescent field showed troublesome for chip-based SMLM acquisition. We found that upon increasing the laser power necessary for SMLM, not only stochastic emission occurred (as desired), but also a spontaneous fluorescent emission remained present through the image acquisition, decreasing the signal-to-background and thus hindering the localization precision of the *d*STORM algorithm. We argue that, while the laser intensity at the waveguide-sample interface was strong enough to induce the photo-switching of the selected marker (namely, the CMDR), the intensity at the tail of the evanescent field reaching deeper into the sample was too weak to enable photo-switching but sufficient enough to support spontaneous fluorescence emission. Also, due to the construct of our optical system, we observed a low photon count of the blinking molecules which further compromised the SMLM reconstruction. We foresee that further efforts could minimize the undesired background signal by a) avoiding unspecific fluorophore binding via waveguide surface functionalization; b) employing advanced labeling techniques such as on-chip point accumulation imaging in nanoscale topography (DNA-PAINT) [2] to ensure low background signal; c) employing thinner tissue samples (~100 nm) to fully exploit the maximum intensities of the evanescent field; and d) modifying the optical setup to improve the photon collection at the camera sensor. In particular, by replacing the current beam splitters with dichroic mirrors that allow a higher transmission rate for the emission spectrum of the fluorescent markers (see *Supplementary Information S11*).

We also encountered initial challenges obtaining consistent labeling repeatability. In particular, we observed significant variability among the staining quality throughout this study, even after following identical labeling protocols across various experiments. We found that the sucrose-methylcellulose droplet (used for the collection of the slices after sectioning) was masking the binding sites on the samples, thus reducing the antigenicity of the targeted proteins. We solved this issue by adjusting the initial washing steps of the cryoprotectant until successful staining was obtained. To this, different temperatures and incubation times were explored, according to existing preparation guidelines for Tokuyasu cryosections [3–6]. Consequently, the samples for chip-based TIRFM, IFON, and SMLM were optimally labeled following incubation in phosphate-buffered saline (PBS) at 37 °C for 30 min, while the samples for chip-based CLEM were successfully stained after incubation in PBS at 0 °C for 20 min (see *Supplementary Information S3*).

Lastly, sectioning artifacts in the form of knife marks, tissue folds, and tissue rupture were also observed along with this study (see *Supplementary Information S7*). These problems were resolved by ensuring optimum blade sharpness on the cryo-ultramicrotome and by adjusting both the sectioning temperature and the slice thickness for each tissue type.

### S6 Photobleaching of membrane markers

To obtain an overall view of tissue sections, a membrane marker is desired. However, membrane probes exhibit a high affinity to the photonic chip surface, resulting in strong absorption of the propagating light along the waveguide. This phenomenon not only limits the excitation intensity reaching the sample but also introduces an undesired background signal that hampers the quality of the chip-TIRFM imaging. To overcome this problem, the power of the excitation beam is temporarily increased to photobleach the fluorescent molecules in the vicinity of the imaging waveguide. Although the emission intensity at the sample location is also reduced, the fluorescent signal of the non-bleached molecules deeper in the sample remains stable through the image acquisition, allowing continuous illumination over prolonged timescales (> 5 min). Arguably, this phenomenon is due to the decaying nature of the evanescent field (see *Supplementary Information S2*). We hypothesize that, since the illumination intensity of the evanescent field is significantly higher at the interface between the coupled waveguide and the sample, the fluorophores in its close vicinity are more susceptible to irreversible photodamage. Further away from the waveguide, the fluorescent markers are exposed to lower excitation intensities and, consequently, less prone to photobleaching. Depending on the coupling efficiency and the geometry of the waveguide, the bleaching process can take few seconds (2 – 10 sec), to around 1 min. Figure S6 shows a 400 μm wide waveguide containing a placental tissue cryosection labeled with CellMask Deep Red. Consecutive frames (1-5) illustrate diverse time points of the bleaching process over a fixed field of view.

**Figure S6.**
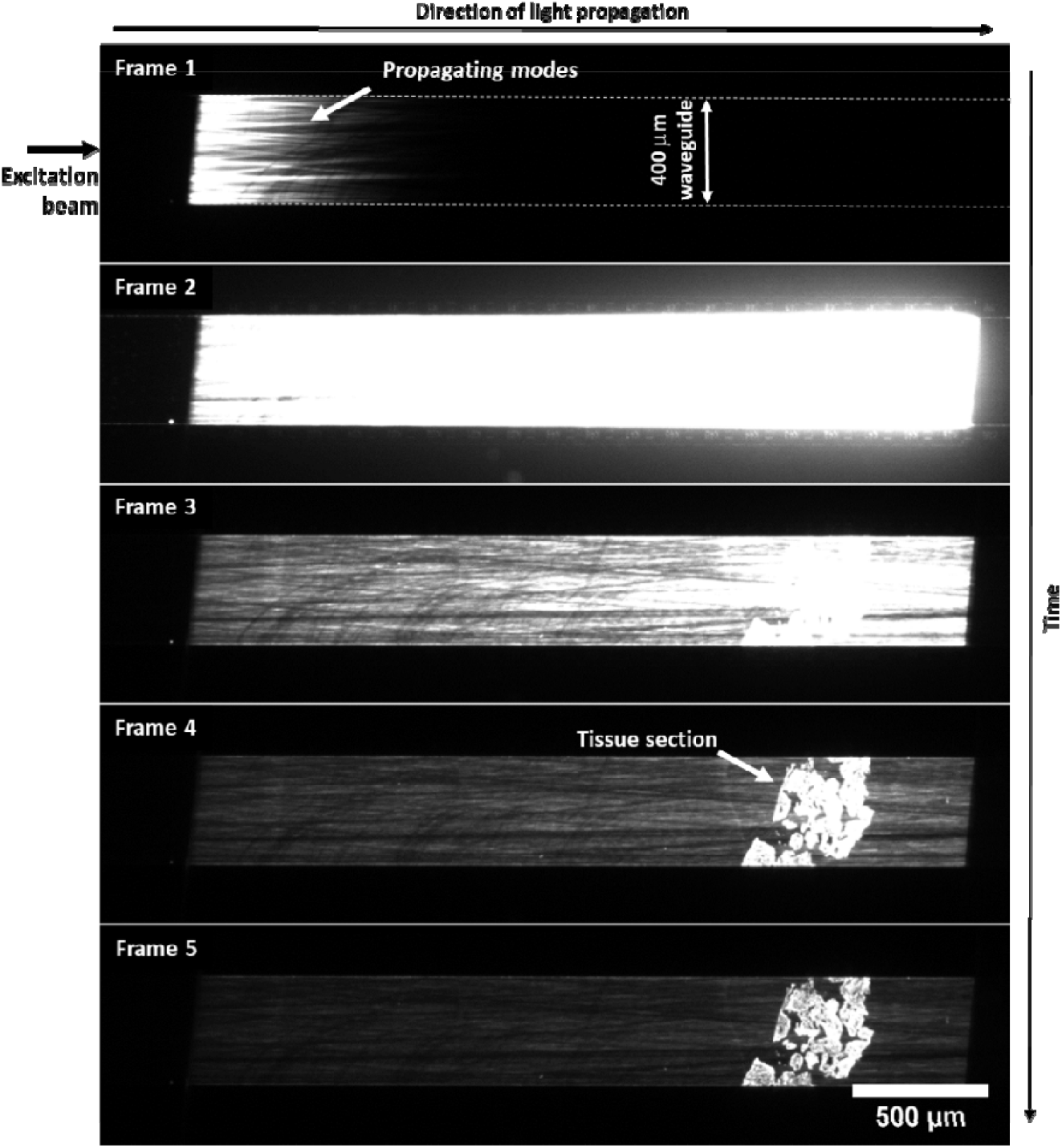
Photobleaching process of a membrane marker bind to a 400 μm waveguide. The power of the excitation beam is temporarily increased to induce photobleaching of the fluorescent molecules in the vicinity of the waveguide (frame 1 to frame 3). After few seconds, the tissue section is revealed (frame 3 to frame 4). Further bleaching dramatically reduces the background signal of the membrane marker, allowing for high-contrast chip-TIRFM imaging (frame 5).

### S7 Sectioning artifacts

To conduct histological analysis, adequate morphological preservation is required. Sectioning artifacts in the form of knife marks (Figure S7a), tissue folds (Figure S7b), and tissue rupture (Figure S7c) are commonly present on the Tokuyasu cryosections. Sectioning parameters such as chamber temperature, slide thickness, and blade sharpness must be carefully adjusted to preserve the structure of the cryosections.

**Figure S7.**
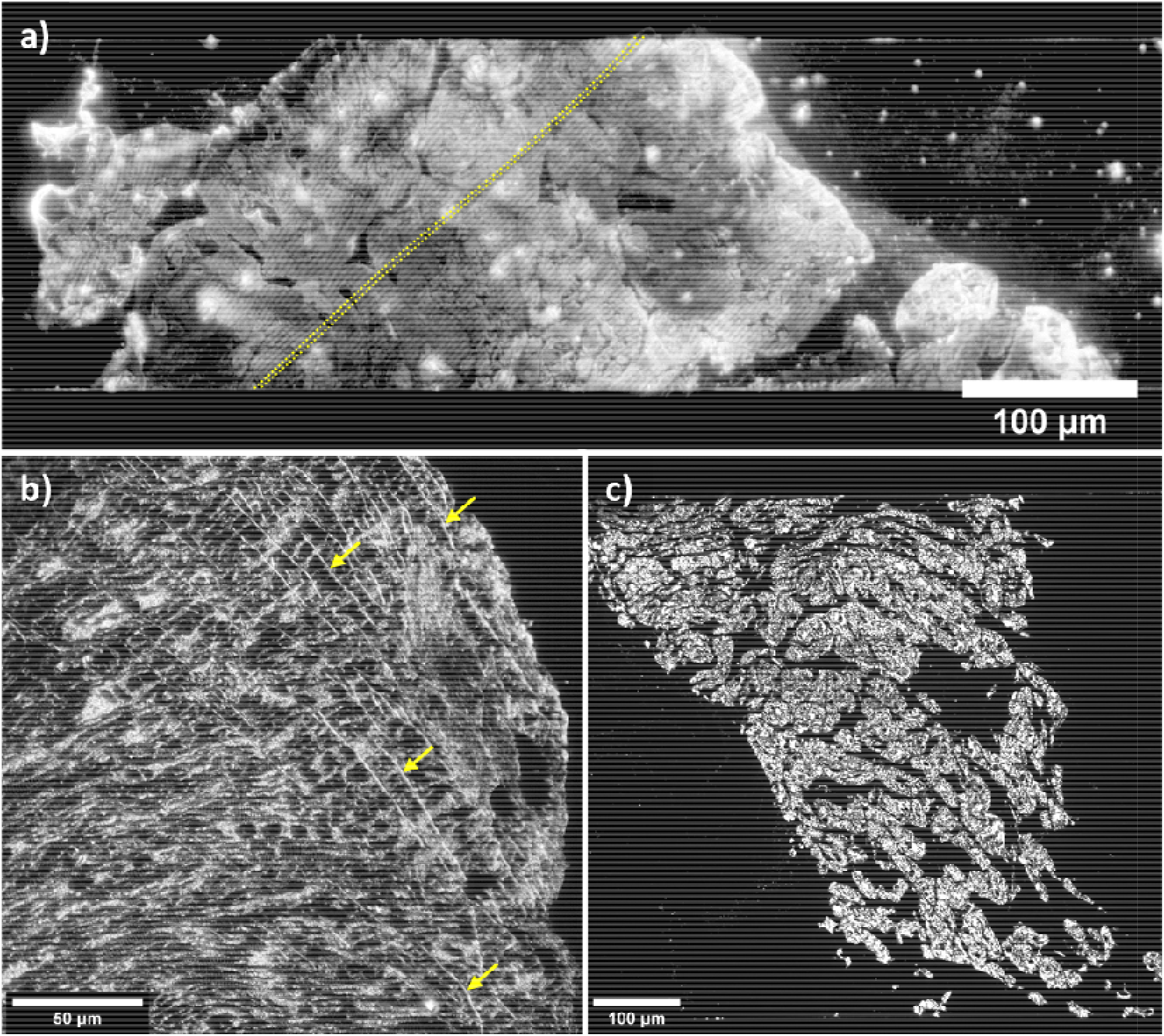
Sectioning artifacts of Tokuyasu cryosections. **(a)** The yellow-dotted lines illustrate the knife marks along a human placental cryosection. **(b)** The yellow arrows denote the location of folds on a pig heart cryosection. **(c)** Illustration of a pig heart tissue sample ruptured during cryosectioning.

### S8 Materials and reagents used for the preparation of Tokuyasu sections

**Table S8a.** Materials and reagents used for the preparation of Tokuyasu sections.

| Material/<br>reagent | Manufacturer | Catalog<br>number | Stock<br>concentration | Working<br>concentration | Purpose |
| --- | --- | --- | --- | --- | --- |
| #1.5 coverslip | VWR | 48393-151 | - | - | Coverslip |
| Picodent twinsil | Picodent | 1300 1000 | - | 1:1 mixture of<br>solution A and<br>B | Dental cement. Gluing and<br>sealing. |
| Poly-L-lysine | Sigma-Aldrich | P8920 | 0.1 % (w/v) in<br>H<sub>2</sub>O | 1:1 | Chip-surface coating for<br>improved adhesion of<br>biological samples |
| CellMask Deep Red<br>(CMDR) | Invitrogen | C10046 | 5 mg/mL | 1:2000 | Membrane staining |
| Phalloidin-Atto565 | Sigma-Aldrich | 94072 | 27.88 mg/mL | 1:100 | F-actin staining |
| Sytox Green | Invitrogen | S7020 | 5 mM | 1:500 | Nuclear staining |
| Ethyl 3-aminobenzoate<br>methanesulfonate<br>(Tricaine) | Sigma-Aldrich | E10521 | 98% | 1:1 | Euthanasia of zebrafish |
| rabbit anti-Tomm20 | Santa Cruz<br>Biotechnology | FL-145 |  | 1:50 | Primary antibody for<br>TOMM20 mitochondrial<br>staining of the zebrafish<br>eye retina |
| AlexaFluor 647<br>AffiniPure F(ab')<sub>2</sub><br>fragment donkey anti<br>rabbit IgG | Jackson<br>Immuno-Research | 711-606-152 |  | 1:200 | Secondary antibody for<br>TOMM20 mitochondrial<br>staining in zebrafish eye<br>retina |
| Texas Red-X<br>Phalloidin | Invitrogen | T7471 |  | 1:50 | F-actin staining of zebrafish<br>eye |
| Podoplanin (hamster<br>anti-mouse)<br>Monoclonal Antibody | ThermoFisher | 14-5381-85 | 0.5 mg/mL | 1:100 | Primary antibody for<br>Podoplanin staining in<br>mouse kidney |
| Goat anti-Hamster<br>IgG (H+L) Alexa Fluor<br>568 | ThermoFisher | A-21112 | 2 mg/mL | 1:250 | Secondary antibody for<br>Podoplanin staining in<br>mouse kidney |
| Phosphate buffered<br>saline (PBS) | Sigma-Aldrich | D8662 | - | 1:1 | Washing steps |
| Prolong Diamond | ThermoFisher | P36961 | - | 1:1 | Antifade mountant |

**Table S8b.**
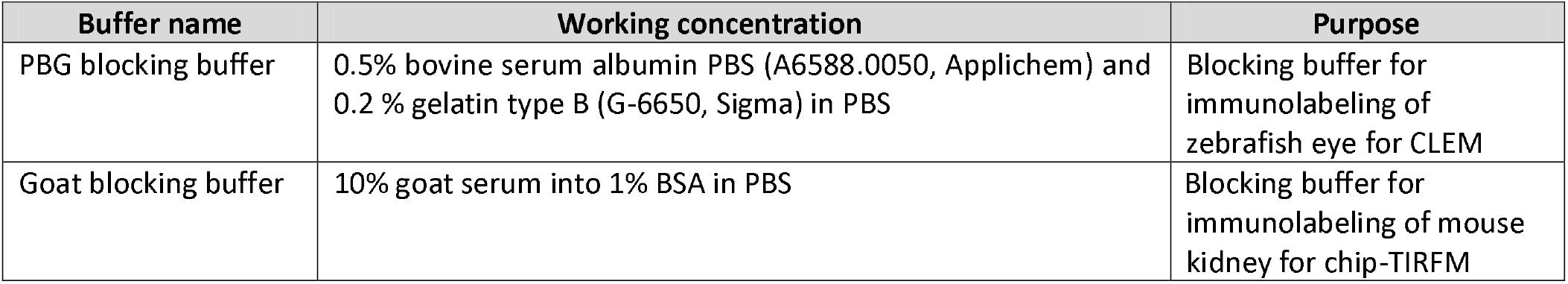
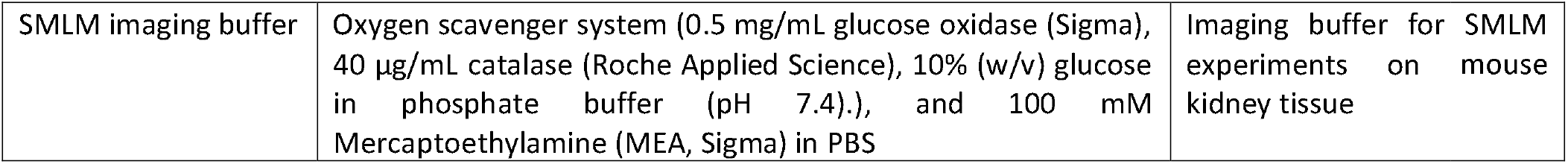
Buffers for sample preparation and imaging of Tokuyasu sections.

### S9 Chip-TIRFM imaging of immunolabeled samples

Fluorescent immunolabeling allows the identification of proteins of interest on the biological samples. The photonic chip not only withstands the chemical and thermal conditions of the sample preparation steps for Tokuyasu cryosections but also allows fluorescent immunolabeling of these samples. Figure S9 shows a 60X magnification image of a 400 nm thick Tokuyasu cryosection of a mouse kidney fluorescently labeled using CellMask Deep Red for membranes (shown in red) and Sytox Green for nuclei (shown in blue). The podoplanin protein was immunolabeled using hamster anti-mouse podoplanin as a primary antibody, and goat anti-hamster conjugated to Alexa Fluor 568 as a secondary antibody (shown in green). *Supplementary Information S8* provides a detailed description of the dyes used for immunolabeling of the mouse kidney cryosection.

**Figure S9.**
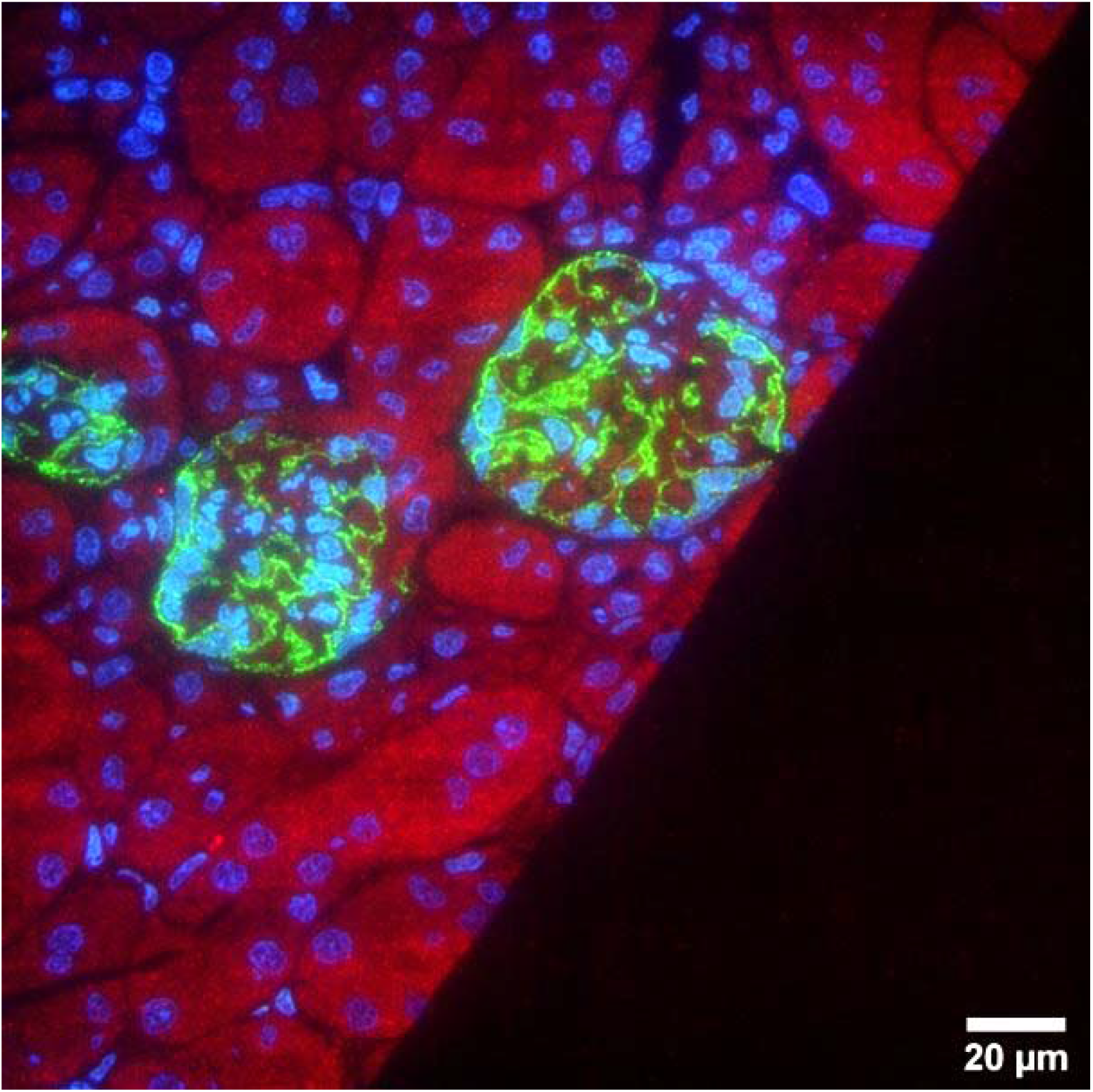
Chip-TIRFM image of a 400 nm thick mouse kidney cryosection fluorescently immunolabeled by Tokuyasu method. Membranes labeled with CellMask Deep Red (pseudo-colored in red) and nuclei labeled with Sytox Green (pseudo-colored in blue). The glomeruli were labeled with hamster anti-mouse podoplanin as a primary antibody, and goat anti-hamster conjugated to Alexa Fluor 568 as a secondary antibody (pseudo-colored in green). The image was collected with a 60X/1.2NA water immersion objective lens.

### S10 Comparative FOV between chip-based IFON and SIM

The photonic chip allows the implementation of advanced microscopy techniques including intensity fluctuation-based optical nanoscopy (IFON) over large fields-of-view (FOV) (Figure S10a). Although structured illumination microscopy (SIM) has been proposed as the fastest super-resolution method for histopathological analyses [7–9], the FOV achieved by this technique is limited to a much smaller area than chip-based IFON when a high magnification objective lens is used. A typical commercial SIM system, e.g., OMX V4 Blaze, GE Healthcare, allows for reconstructed 3D-SIM images of approximately 40 × 40 μm^2^ using a 60X/1.42NA oil immersion objective. Hence, to achieve a similar FOV to that of chip-based IFON, a tile mosaic composed of 7 × 7 reconstructed 3D-SIM images is needed (Figure S10b). Considering that a set of 15 raw images are required for each of the 8 z-planes necessaries to reconstruct a single SIM image, and accounting for the 10 μm overlap between adjacent images (Figure S10b), a total of 5880 SIM raw images are needed for an equivalent FOV as the one obtained with the photonic chip (15 raw images/z-plane × 8 z-planes/3D-SIM × 49 3D-SIM = 5880 raw images). Also, considering a typical image acquisition of 30 msec per raw image and a reconstruction time of 3 min per 3D-SIM image rounds up to a total imaging time of 2.5h from acquisition (30 msec/raw image × 5880 raw images = 176.4 sec 3 min) to 3D-SIM reconstruction (3 min/3D-SIM image × 49 3D-SIM images = 147 min). Importantly, we achieved a high-resolution chip-based IFON image over a fixed FOV of 220 × 220 μm^2^ after collecting a relatively small image stack of 500-frames using a 60X/1.2 water immersion objective, requiring approximately 10 min from acquisition to image reconstruction. We acknowledge that the implementation of a 2D-SIM scheme reduces the amount of acquired data (e.g., 9 raw images/2D-SIM × 49 2D-SIM = 441 raw images) and, consequently improves the processing time for a single-plane 2D-SIM, potentially becoming faster than chip-based IFON. We could not benchmark the exact numbers for this premise, since the SIM microscope available at our facilities only allows for 3D-SIM, and requires a z-stack of at least 7 to 8 planes to properly reconstruct an imaging area of 40 × 40 μm^2^ . Nevertheless, the chip-based method offers a much less complex and highly cost-effective alternative to a commercial SIM instrument.

**Figure S10.**
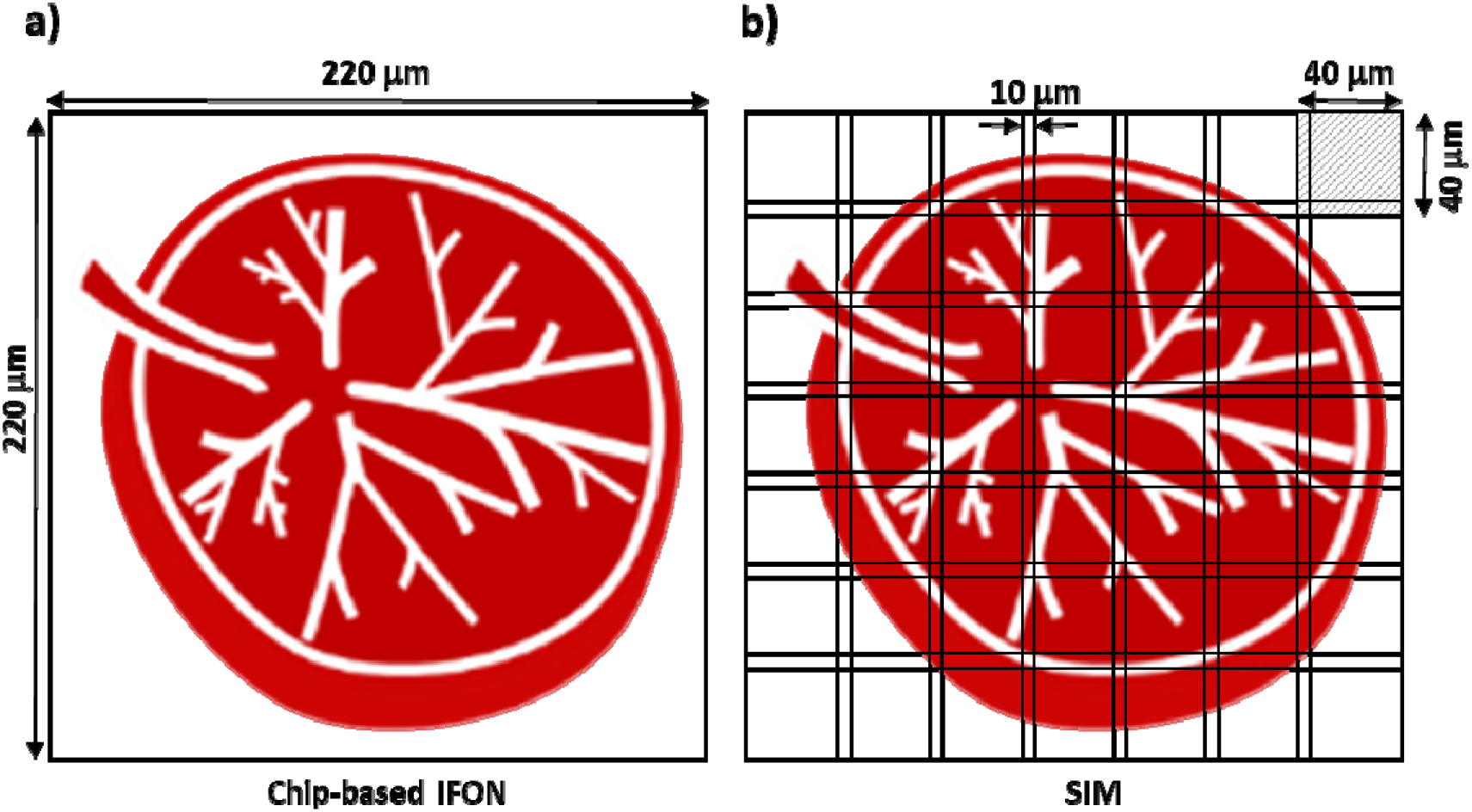
Comparative FOV between chip-based IFON and 3D-SIM. **(a)** Chip-based IFON allows a FOV of 220 × 220 μm^2^ after collection of an image stack of 500-frames and reconstruction time of approx. 10 min. **(b)** To achieve a similar FOV with 3D-SIM, a tile mosaic image is constructed. It requires the acquisition of 5880 raw images and a reconstruction time of 2.5h. The upper-right square denotes the typical FOV attainable with 3D-SIM (40 × 40 μm^2^).

### S11 Detailed description of the chip-TIRFM setup

The chip-TIRFM setup is composed of two main parts, namely the collection path and a photonic chip module, as illustrated in Figure S11a. The collection path consists of a commercial upright microscope equipped with an emission filter set (see Table S11), a sCMOS camera, and conventional microscope objective lenses of diverse magnifications, which can be interchanged depending on the imaging needs. Figures S11b and S11c provide a detailed view of the chip-TIRFM setup.

**Table S11.** Longpass and bandpass filters used in the setup for image acquisition

| Excitation wavelength (nm) | Emission filter set |  |
| --- | --- | --- |
|  | Long-pass filter (nm) | Band-pass filter (nm) |
| 488 | 488 | 520/36 |
| 561 | 561 | 591/43 |
| 640 | 664 | 690/40 |

**Figure S11.**
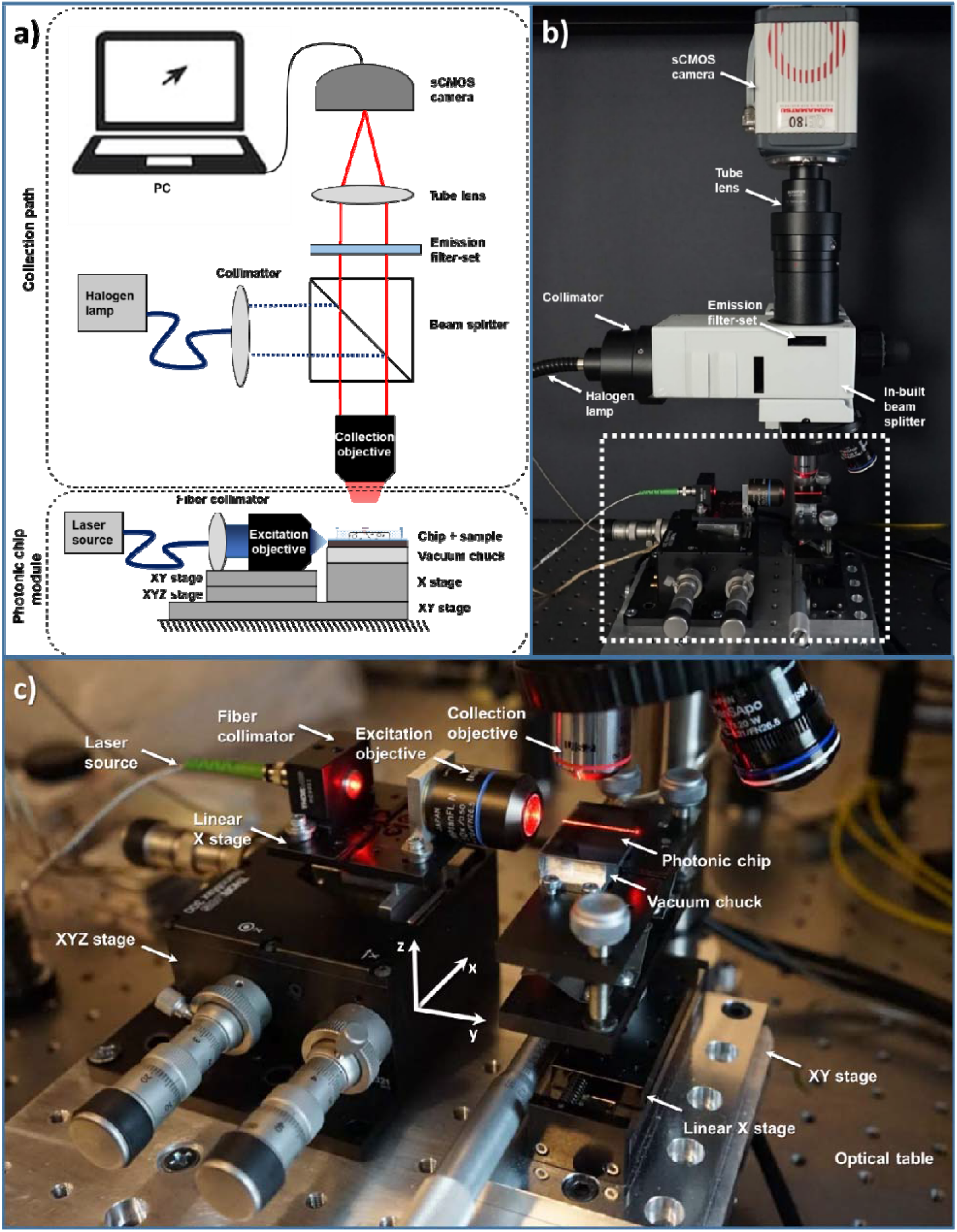
Chip-TIRFM setup. **(a)** Schematic representation of the chip-TIRFM setup illustrating the collection path and the photonic chip module. **(b)** Side view of a chip-TIRFM setup denoting the collection path components. The dotted-white box represents the photonic chip module shown in (c). **(c)** Close view of the photonic chip module components.

### S12 SEM imaging on a photonic chip

For correlative light-electron microscopy (CLEM), the Tokuyasu cryosections are imaged on a scanning electron microscope (SEM) after completion of chip-TIRFM imaging. To this, the coverslip and the PDMS frame are removed. Then, the sample is post-fixed with 0.1% glutaraldehyde, masked with methylcellulose, and further coated with a 10 nm layer of platinum/carbon. The photonic chip is placed on a 25 mm Pin Mount (Figure S12A) and transferred to a SEM. A bright-field image assists in finding the sample (Figure S12B). A low accelerating voltage allows high-resolution SEM imaging of the sample (Figure S12C).

**Figure S12.**
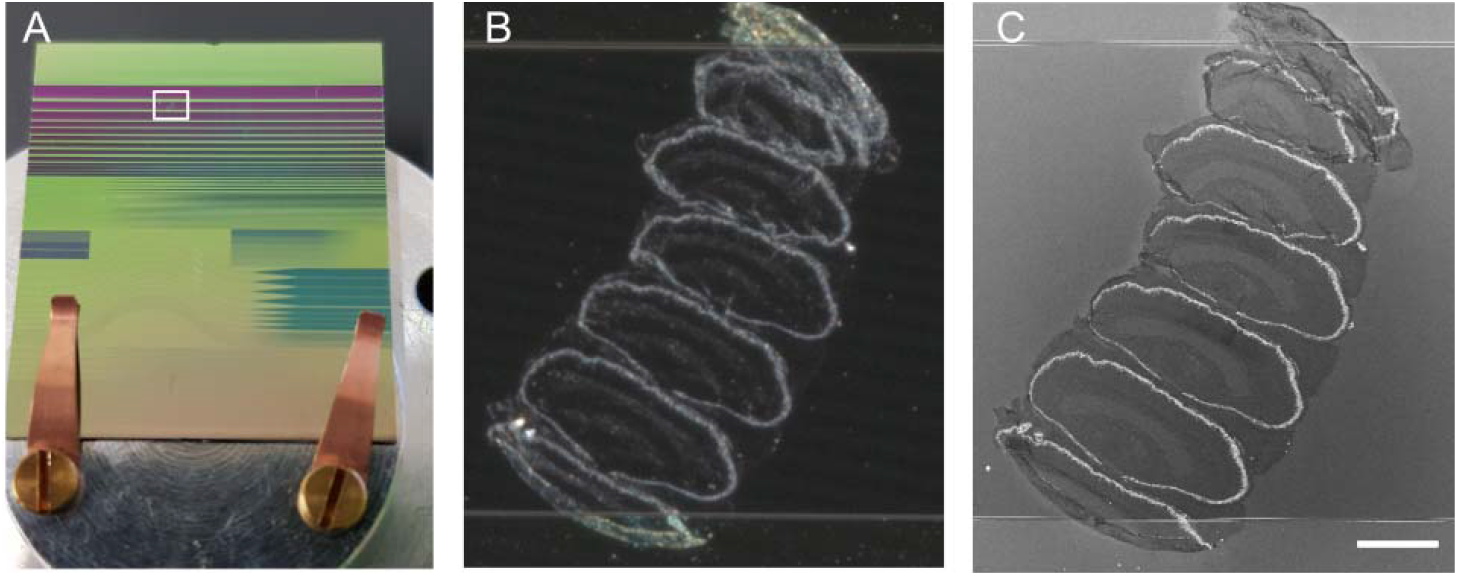
Chip-based CLEM imaging. **(A)** Photonic chip mounted on a 25 mm pin stub for imaging in the SEM. The white frame corresponds to the zoomed area in (B) and (C). **(B)** Bright-field image of zebrafish retina serial sections on a 600 μm strip waveguide. **(C)** The same area acquired with SEM. Scale bar 100 μm.

